# Interaction of two strongly divergent archaellins stabilizes the structure of the *Halorubrum* archaellum

**DOI:** 10.1101/836379

**Authors:** Mikhail G. Pyatibratov, Alexey S. Syutkin, Tessa E.F. Quax, Tatjana N. Melnik, R. Thane Papke, Johann Peter Gogarten, Igor I. Kireev, Alexey K. Surin, Sergei N. Beznosov, Anna V. Galeva, Oleg V. Fedorov

**Affiliations:** Institute of Protein Research, Russian Academy of Sciences, Institutskaya st. 4, Pushchino, Moscow Region, 142290, Russia; University of Freiburg, Institute for Biology II- Microbiology, Archaeal virus-host interactions, Schänzlestrasse 1, 79104 Freiburg, Germany; Department of Molecular and Cell Biology, University of Connecticut, Storrs, CT 06269-3125, USA; A.N. Belozersky Institute of Physico-chemical Biology, M.V. Lomonosov Moscow State University, Leninskie Gori 1, Bldg 40, Moscow 119234, Russia; Pushchino Branch, Shemyakin–Ovchinnikov Institute of Bioorganic Chemistry, Russian Academy of Sciences, Prospekt Nauki 6, Pushchino, Moscow Region, 142290, Russia; State Research Center for Applied Microbiology & Biotechnology, Obolensk, Serpukhov district, Moscow Region, 142279, Russia

## Abstract

The archaellum is a unique motility structure that has only functional similarity to its bacterial counterpart, the flagellum. Archaellar filaments consist of thousands of copies of the protein protomer archaellin. Most euryarchaeal genomes encode multiple homologous archaellins. The role of these multiple archaellin genes remains unclear. Halophilic archaea from the genus *Halorubrum* possess two archaellin genes, *flaB1* and *flaB2*. Amino acid sequences of the corresponding protein products are extraordinarily diverged (identity of ∼ 40%). To clarify roles for each archaellin, we compared archaella from two natural *Halorubrum lacusprofundi* strains: the DL18 strain, which possesses both archaellin genes, and the type strain ACAM 34 whose genome contains the *flaB2* gene only. Both strains synthesize functional archaella; however, the DL18 strain, where both archaellins are present in comparable amounts, is more motile. In addition, we expressed these different *Hrr. lacusprofundi* archaellins in a *Haloferax volcanii* strain from which the endogenous archaellin genes were deleted. Three *Hfx. volcanii* strains expressing *Hrr. lacusprofundi* archaellins *flaB1*, *flaB2* or *flaB1-flaB2* produced archaellum filaments consisting of only one (FlaB1 or FlaB2) or both (FlaB1/B2) archaellins. All three recombinant *Hfx. volcanii* strains were motile, although there were profound differences in the efficiency of motility. The recombinant filaments resemble the natural filaments of *Hrr. lacusprofundi*. Electron microscopy showed that FlaB1 FlaB2-archaella look like typical supercoiled filaments, while with the shape of the FlaB1- and FlaB2-archaella is more variable. Both native and recombinant FlaB1 FlaB2-filaments have greater thermal stability and are more resistant to low salinity stress than single-component filaments. This shows that thermal stability of archaellins depends on the presence of both archaellin types, indicating a close interaction between these subunits in the supramolecular structure. Functional helical *Hrr. lacusprofundi* archaella can be composed of either single archaellin: FlaB2 or FlaB1; however, the two divergent archaellin subunits in combination provide additional stabilization to the archaellum structure and thus adaptation to a wider range of external conditions. A comparative genomic analysis of archaellins suggests that the described combination of divergent archaellins is not restricted to *Hrr. lacusprofundi,* but is occurring also in organisms from other haloarchaeal genera.

## INTRODUCTION

Archaeal flagella (archaella) are morphologically and functionally similar to bacterial flagella. However, the archaellum structure, assembly mechanism and protein composition is fundamentally different from the flagellum and instead shows similarity to type IV pili. Fundamental differences between archaeal and bacterial flagellar filaments are the presence of signal sequences in archaeal flagellins (archaellins) and the multiplicity of archaellin encoding genes in archaea (Jarrell and Albers, 2012). Recently, cryoelectron microscopy was used to determine the spatial structures of the archaella filaments of three archaea: two methanogenes *Methanospirillum hungatei* (3.4 Å resolution) (Poweleit *et al.*, 2016) and *Methanococcus maripaludis* (4 Å resolution) (Meshcheryakov *et al.*, 2019) and a hyperthermophile *Pyrococcus furiosus* (4.2 Å resolution) (Daum *et al.*, 2017). Furthermore the crystal structure of the N-terminal truncation of archaellin FlaB1 *Methanocaldococcus jannaschii* has been determined at a resolution of 1.5 Å (Meshcheryakov *et al*., 2019). The structure of archaeal filaments differs significantly not only from bacterial flagella, but also from bacterial type IV pili (Braun *et al*., 2016; Poweleit *et al*., 2016). The amino acid residues of archaellins responsible for intersubunit interactions were identified, as well as the protein regions forming the outside surface of the filament. The proposed models for the archaellar filament do not contain the long-pitch protofilaments found in bacterial flagellar filaments. To explain the archaella supercoiling, it was proposed to consider them as semiflexible filaments in a viscous medium (Wolgemuth *et al.*, 2000; Tony *et al.*, 2006; Coq *et al.*, 2008). For such structures, thrust can be generated by their rotation. Using molecular modeling, it was shown that conformational changes in the globular domain of the archaellin can lead to extension and compression, as well as bending of the filaments (Braun et al., 2016). However, the detailed mechanism of the archaellum supercoiling is not fully understood.

The proposed models of spatial archaellar filament structure do not explain the structural and functional role of multiple archaellins. Despite the presence of several archaellin genes in genomes of *M. hungatei*, *M. maripaludis* and *P. furiosus*, protein products of only one of these genes were found incorporated in their filaments (Poweleit *et al.*, 2016; Daum *et al.*, 2017; Meshcheryakov *et al.*, 2019). This raises the question about the importance of encoding multiple different archaellins. Interestingly, the presence of several copies of archaellin genes in archaeal genomes is very common.

Currently, almost 3,000 archaeal genomic sequences are deposited at the NCBI database (https://www.ncbi.nlm.nih.gov/genome/browse/#!/prokaryotes/archaea), including about 400 genomes of halophilic archaea. The majority of these archaeal genomes contain archaellin genes. The known genomes of crenarchaeota typically only possess a single archaellin gene. However, the large majority of euryarchaeal genomes contain multiple archaellin genes.

In haloarchaea, multiple archaellin genes appear to have originated from duplication events (Desmond *et al.*, 2007). Duplicated genes can be located in the same or different operons. For example, in three popular model haloarchaea the organization of archaellin genes differ significantly. *Haloarcula hispanica* has three operons with 1 archaellin gene in each of them, two *Haloferax volcanii* archaellin genes are in one operon, and *Halobacterium salinarum* has two operons with 2 and 3 archaellin genes each. The similarity between the archaellin paralogs of the above species is still very high, indicating relatively recent duplication. Interestingly, several haloarchaea have multiple archaellin genes (most often, two) that are very divergent (such as *Halobiforma*, *Halopiger*, *Halorubrum*, *Natrialba* and *Natronolimnobius* species). Even archaea belonging to the same genus can differ drastically from each other in the number and size of the archaellin genes. It has been suggested that the archaellum helicity, as for bacterial flagella, can be achieved through a combination of subfilaments of different lengths, constructed from different types of subunits (Tarasov *et al.*, 2000; 2004). For *Hbt. salinarum* it was demonstrated that both archaellins FlgA1 and FlgA2 are necessary for the formation of a functional helical archaellum, and mutant strains with a single FlgA1 or FlgA2 archaellin had straight non-functional filaments. In the case of methanogenic archaea, the multiple archaellin genes were shown to encode major and minor structural components of the archaellum filament. A ‘hook’-like structure was observed in a number of methanogenic archaea. In *Methanococcus voltae* and *M. maripaludis* the archaellum hook segment is built of the FlaB3 archaellin, while the FlaB1 and FlaB2 proteins are the main components of the filament (Bardy *et al.*, 2002; Chaban *et al.*, 2007). Inactivation of either the *flaB1* or *flaB2* genes resulted in loss of motility and cessation of archaellum synthesis (including the hook) (Chaban *et al.*, 2007). Recently, it has been shown that FlaB1 is the predominant component of *Methanococcus maripaludis* filaments (Meshcheryakov *et al.*, 2019). Inactivation of the *flaB3* gene does not lead to the cessation of filament synthesis and a noticeable change in motility on semi-liquid agar (Chaban et al., 2007). However, time-lapse microscopy showed impaired motility for this deletion strain (movement in a closed circle). The structures corresponding to the *Methanococcales* hooks were not found in the native archaella of other archaea. It is possible, that the differentiation of one of the archaellins into a “hook protein” with a special structural role is a relatively late evolutionary event in the *Methanococcales* and is not typical for other archaea.

In *Hfx. volcanii* it was shown that the archaellum filament consists of one major (FlgA1) and one minor (FlgA2) component. However, the structural role of the minor component is unknown and it does not form a hook-like structure (Tripepi *et al.*, 2013). Deletion of the *flgA2* gene leads to hypermotile cells by an unknown mechanism (Tripepi *et al.*, 2013).

In contrast to all above mentioned examples, some halophilic archaea can form functional archaella with only one of the genomically encoded archaellin proteins (Pyatibratov *et al.*, 2008; Syutkin *et al.*, 2012, 2014). These archaellins were thought to be partly redundant. However, in the case of *Haloarcula marismortui* it was shown that these proteins function as ecoparalogs, i.e., they are expressed under different environmental conditions, and provide distinct stability advantages under varying salt concentrations (Syutkin *et al.*, 2014; 2019).

In this work we investigate the role of multiple archaellin genes of the *Halorubrum* genus. In contrast to the systems studied before, members of the *Halorubrum* group, possess multiple archaellins with highly diverged protein sequences. Our preceding work has shown that functional helical archaella filaments of *Hrr. lacusprofundi* ATCC49239 (ACAM 34) are formed from a protein encoded by a single archaellin gene (*flaB2*) (Syutkin *et al.*, 2012). However, unlike *Hrr. lacusprofundi* ACAM 34, other *Halorubrum* species possess at least two archaellin genes (*flaB1* and *flaB2)* located in one operon. The amino acid sequences of the FlaB1 and FlaB2 archaellins differ significantly from each other (<43% identical residues, the N-terminal region being more conserved). We use the *Hrr. lacusprofundi* archaella as a model to address the role of multiple highly divergent archaellin genes often found in genomes of haloarchaea.

Recently *Hrr. lacusprofundi* strains (DL18 and R1S1) with two archaellin genes (which is more typical for *Halorubrum* species) were isolated (Tschitschko *et al.*, 2018). Since the presence of a single archaellin gene in *Hrr. lacusprofundi* ACAM 34 is sufficient to form functional helical archaella, the presence of the *flaB1* gene may seem redundant. FlaB1 could be responsible for: (i) formation of specific filaments that differ from FlaB2 in function (and for example function as ecoparalogs); (ii) stabilization of the filament structure together with FlaB, possibly as a result of constructive neutral evolution (Lukeš *et al.*, 2011).

In the present study, we characterized the archaella of the *Hrr. lacusprofundi* DL18 strain, containing two archaellin genes *flaB1* and *flaB2,* and compared them with archaella of the ACAM 34 strain whose FlaB2 is (with the exception of the signal sequence) completely identical to its counterpart from the DL18 strain. To clarify the role of FlaB1 and FlaB2 archaellins, we used the well-developed expression system of *Hfx. volcanii*. With this approach we could show that either the FlaB1 or FlaB2 protein is sufficient to form functional archaella. However, the combination of the two proteins renders the archaellum filament structure much more stable and consequently leads to the highest motility.

## RESULTS

### Comparison of native FlaB2 and FlaB1/FlaB2 filaments

First we analyzed the archaellin amino acid sequences of both *Hrr. lacusprofundi* strains. Remarkably, this analysis showed that the signal peptide MFETILNEEERG encoded by the ACAM 34 *flaB2* gene is different from the DL18 FlaB2 signal peptide MFEFINNDKDRG and identical to that of DL18-FlaB1 (**Figure S1**). At the same time, the nucleotide sequences of the *flaB2* genes of both strains, with the exception of the region encoding the signal peptide, are completely identical (**Figure S2**). Thus, the disappearance of the *flaB1* gene in the ACAM 34 strain can be explained by a recombination and subsequent deletion event of a DNA segment in an ancestral two-archaellin operon.

*Hrr. lacusprofundi* DL18 FlaB1 and FlaB2 are strongly diverged (identity 36%), as in other *Halorubrum* species (**Figure S1**). This strikingly distinguishes them from other haloarchaeal systems. For example, the identity between the two *Hfx. volcanii* archaellins (FlgA1 and FlgA2) is 60%, between those of *Har. hispanica* (FlgA1, FlgB, FlgA2) it is at least 55%, and in *Hbt. salinarum* (FlgA1, FlgA2, FlgB1-FlgB3) the identity is over 80%. *Hrr. lacusprofundi* DL18 FlaB1 and FlaB2 also differ significantly in molecular weight: 19796.67 and 23593.45, respectively.

Both *Hrr. lacusprofundi* strains were isolated from the relict hypersaline Deep Lake in Antarctica (Franzmann *et al.*, 1988; Liao *et al.*, 2016). Due to its high salinity, this lake never freezes and its surface temperature ranges from ⎯20 °C to +10 °C depending on the season. In the laboratory *Hrr. lacusprofundi* cells are able to grow at temperatures ranging from –1 °C to +44 °C (optimum temperature is 33 °C) (Franzmann *et al.*, 1988).

Earlier, we showed that the cells of the *Hrr. lacusprofundi* ACAM 34 strain are motile on semi-liquid media (Syutkin *et al.*, 2012). We compared the motility of both strains on semi-liquid 0.19% agar under the same conditions and found that the DL18 strain shows significantly higher motility (Figure 1). It should be noted that at the same time, the growth rate of the DL18 is higher than that of the ACAM 34. The maximum archaella yield in the late stationary phase was approximately 10 mg per 1 liter culture for both strains. In contrast to the ACAM 34, the cells of the DL18 strain demonstrate a more stable motility and archaella production. To obtain the relatively motile ACAM 34 cells with high archaella yield that were used in the above experiment, it was necessary to pass cells through semi-liquid (0.19%) agar with 2-3 cycles of selection of the most motile cells (Syutkin *et al.*, 2012). When ACAM 34 cells that were kept for a long time (about 1 month) in a liquid medium, were used to inoculate, the archaella yield decreases dramatically.

**Figure 1.**
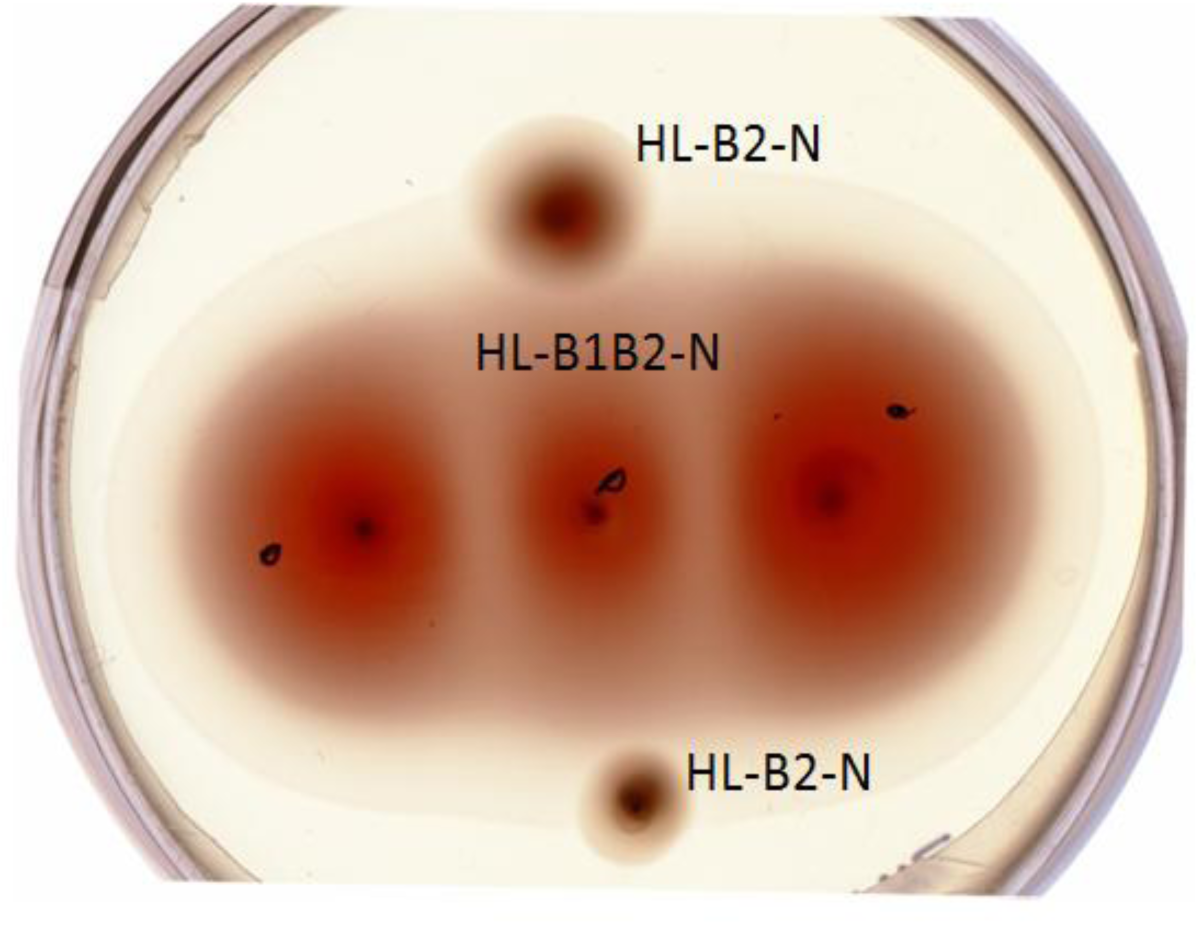
Comparison of cell motility of *Hrr. lacusprofundi* ACAM 34 (HL-B2-N) (in the top and bottom spots) and DL18 (HL-B1B2-N) strains (in three central spots), St-HL medium, 0.19% agar, 25 °C, 28 days.

SDS-PAGE of *Hrr. lacusprofundi* ACAM 34 archaellum filaments (Figure 2) showed a single major band, corresponding to a molecular mass of ∼50 kDa. For the DL18 strain, the same and additional major bands of ∼37 kDa were observed (Figure 2). Mass spectrometry analysis (**Figure S3**) confirmed that the isolated proteins were *Hrr. lacusprofundi* archaellins FlaB2 (50 kDa) and FlaB1 (37 kDa). The apparent molecular masses determined by SDS gel electrophoresis are higher than the true values (23.6 kDa and 19.8 kDa for FlaB1 and FlaB2 respectively), which is typical for halophilic archaellins due to high content of carbonic acids and posttranslational modifications (Gerl and Sumper, 1988; Fedorov *et al.*, 1994; Pyatibratov *et al.*, 2008). The *Hrr. lacusprofundi* DL18 archaellum consists of both FlaB1 and FlaB2 archaellins, present in comparable quantities. The molar ratio FlaB1/FlaB2, determined by measuring the intensity of the protein bands on SDS-PAGE gels, is approximately 1:1 (0.90±0.04) (Figure 2). The archaella, isolated from natural DL18 and ACAM 34 strains, were designated as HL-B1B2-N and HL-B2-N respectively.

**Figure 2.**
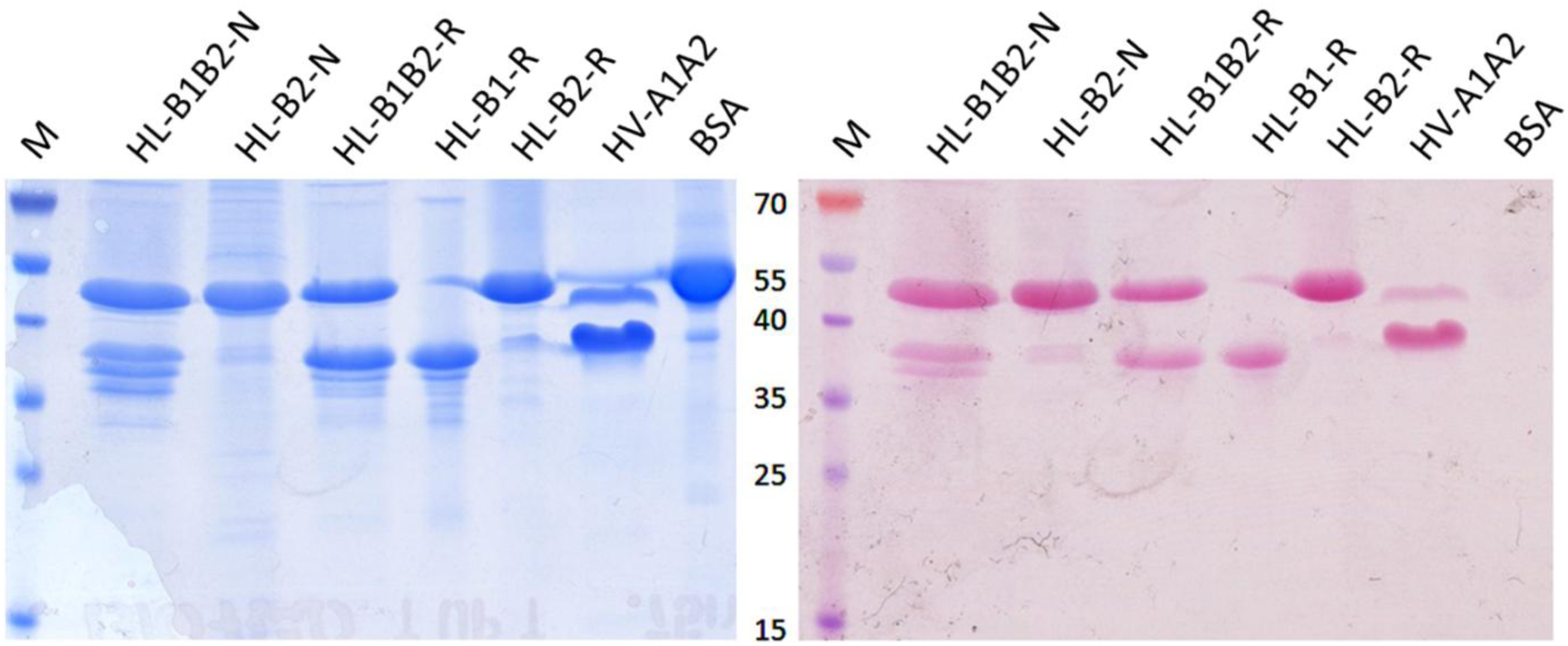
Archaellin staining with Schiff’s reagent in 12.5% polyacrylamide gel (right); the same gel stained with Coomassie G250 (left). M - prestained protein standard, HL-B1B2-N and HL-B2-N – natural archaella of *Hrr. lacusprofundi* DL18 and ACAM 34, HL-B1B2-R, HL-B1-R, HL-B2-R – recombinant archaella isolated from *Hfx. volcanii* MT2 transformed with appropriate plasmids: HV-A1A2 – *Hfx. volcanii* MT45 archaella (positive control), BSA - bovine serum albumin (negative control).

The archaella isolated from the natural DL18 and ACAM 34 strains were observed by electron microscopy, which showed that the HL-B2-N archaella are quite flexible and often twisted in loops and tangles (Figure 3). The HL-B1B2-N-archaella, on the other hand, were generally longer and less flexible (Figure 3). The thickness of the HL-B1B2-N and the HL-B2-N archaella were both ∼10 nm. No structures resembling hooks or basal bodies were observed. The HL-B1B2-N-filaments were homogeneous; no filaments twisted into loops characteristic of the HL-B2-N archaella were observed. This suggests that all isolated archaella from DL18 are composed of a mixture of FlaB1 and FlaB2 archaellins, and that the DL18 strain does not produce two types of filaments made of two different archaellins.

**Figure 3.**
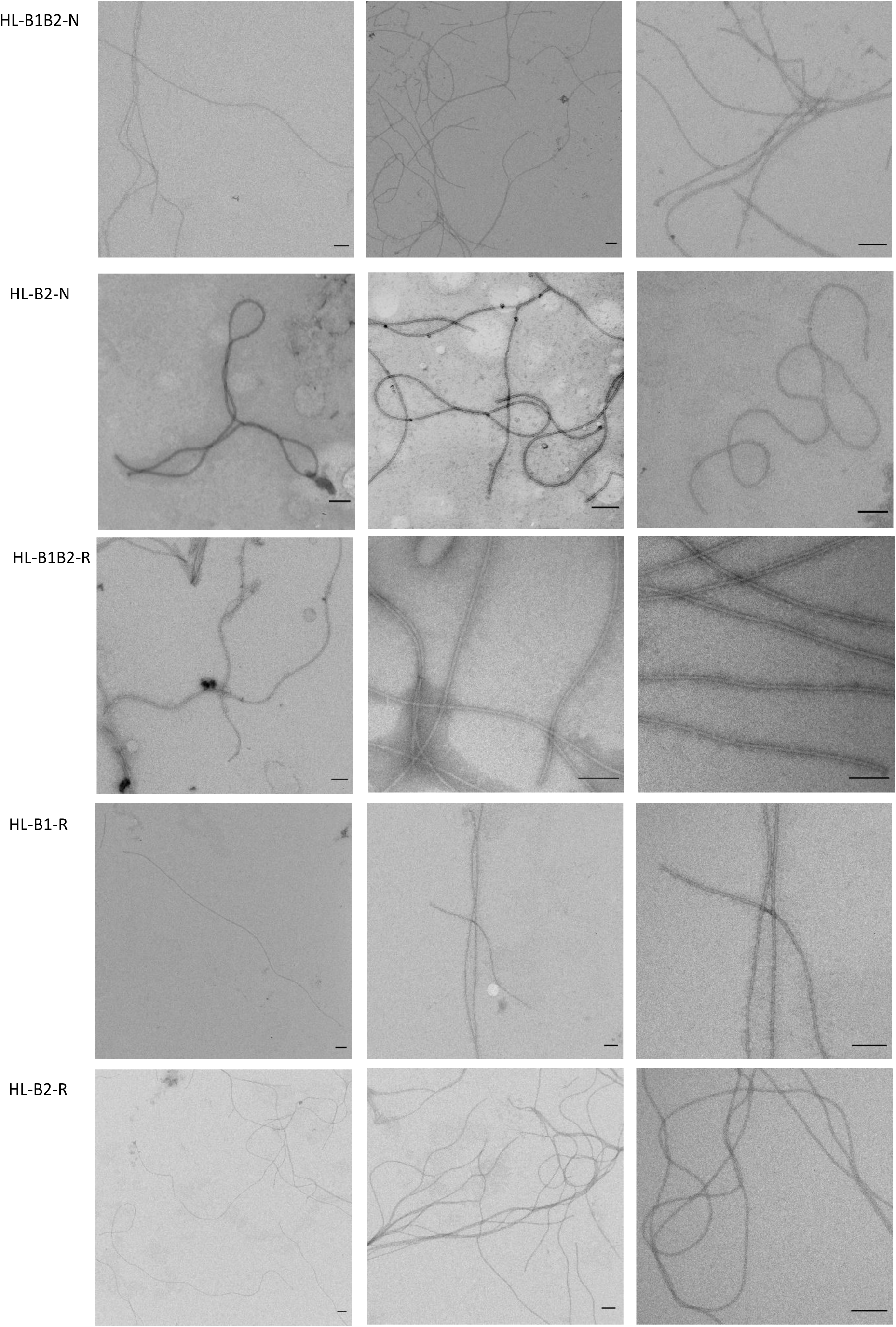
Negatively stained (1% uranyl acetate) preparations of *Hrr. lacusprofundi* ACAM 34 and DL18 archaellar filaments (HL-B2-N and HL-B1B2-N, respectively) and preparations of recombinant *Hrr. lacusprofundi* archaellar filaments HL-B1B2-R, HL-B1-R and HL-B2-R isolated from *Hfx. volcanii* MT2 in 20% NaCl, 10 mM Na-phosphate, pH 8.0. Scale bar – 100 nm.

### Heterologous expression of *Halorubrum* archaellins in *Haloferax volcanii*

The analysis of natural *Halorubrum* strains allowed us to isolate archaellar filaments consisting of FlaB1/FlaB2 and FlaB2 archaellins. Next, we aimed to study if the FlaB1 archaellin is also capable of producing functional archaella. To this end we expressed the different archaellin genes in the heterologous host *Hfx. volcanii* MT2 in which the *flgA1flgA2* archaellin operon was removed (Tripepi *et al.*, 2010). The *flaB1* and *flaB2* genes (separately and together as an operon) were cloned into corresponding plasmids based on the pTA1228 vector (*Amp^r^*, *pyrE2* and *hdrB* markers, inducible ptna promoter) (Allers *et al.*, 2010). After transformation, *Hfx. volcanii* cells were grown in the presence of tryptophan as an inducer of the archaellin expression. Synthesis of recombinant archaella was confirmed by mass spectrometry (**Figure S12-S14**) and electron microscopy (Figure 3**, S4**). Expression of *Hrr. lacusprofundi* archaellins in non-motile *Hfx. volcanii* Δ*flgA1flgA2* leads to the restoration of motility (Figures 4**, S3**). Motility was demonstrated by cells with each of the three archaella types (HL-B1-R, HL-B2-R, HL-B1B2-R, R - recombinant). The HL-B1B2-R strain had the best motility on semi-liquid agar (measured by the diameter of the motility ring) compared with the HL-B1-R and HL-B2-R strains. At the same time, the motility on semi-liquid agar in all three strains was markedly less than in *Hfx. volcanii* strain expressing their natural archaellins (FlgA1/FlgA2) (Figures 4**, S3**). Electron microscopy confirmed the presence of archaella bundles in HL-B1-R, HL-B2-R and HL-B1B2-R cells. HL-B1-R and HL-B1B2-R cells were noticeably more archaellated than HL-B2-R cells, for which specimens without archaella were often observed (**Figure S4**). In addition, we analyzed the swimming behavior of the three strains in liquid medium with live microscopy and compared this with a *Hfx. volcanii* strain with a pMT21 plasmid containing *flgA1flgA2His* (Tripepi *et al*., 2010) that synthesizes the functional archaella when cells are grown in the presence of tryptophan. There were no significant differences in the velocity or the frequency of reversals between any of the strains (**Figure S3).** However, in the *flaB2* expressing strain, an extremely low percentage of cells was motile (<5%) (Figure 5**, Movie S1).** This is in contrast with the other strains, for which a large fraction of cells is motile in liquid medium in early exponential phase **(Movie S2-S4**). Expression of *flaB1* alone results in motile cells, corresponding with the analysis on semi-solid agar plate. However, in liquid medium, the percentage of motile cells is significantly lower (∼30%) than for the *flaB1flaB2* expression strain (∼60%) (Figure 5). Thus, the expression of FlaB1 only archaellum filaments might lead to slightly less motile cells in comparison to the two-archaellin filaments.

**Figure 4.**
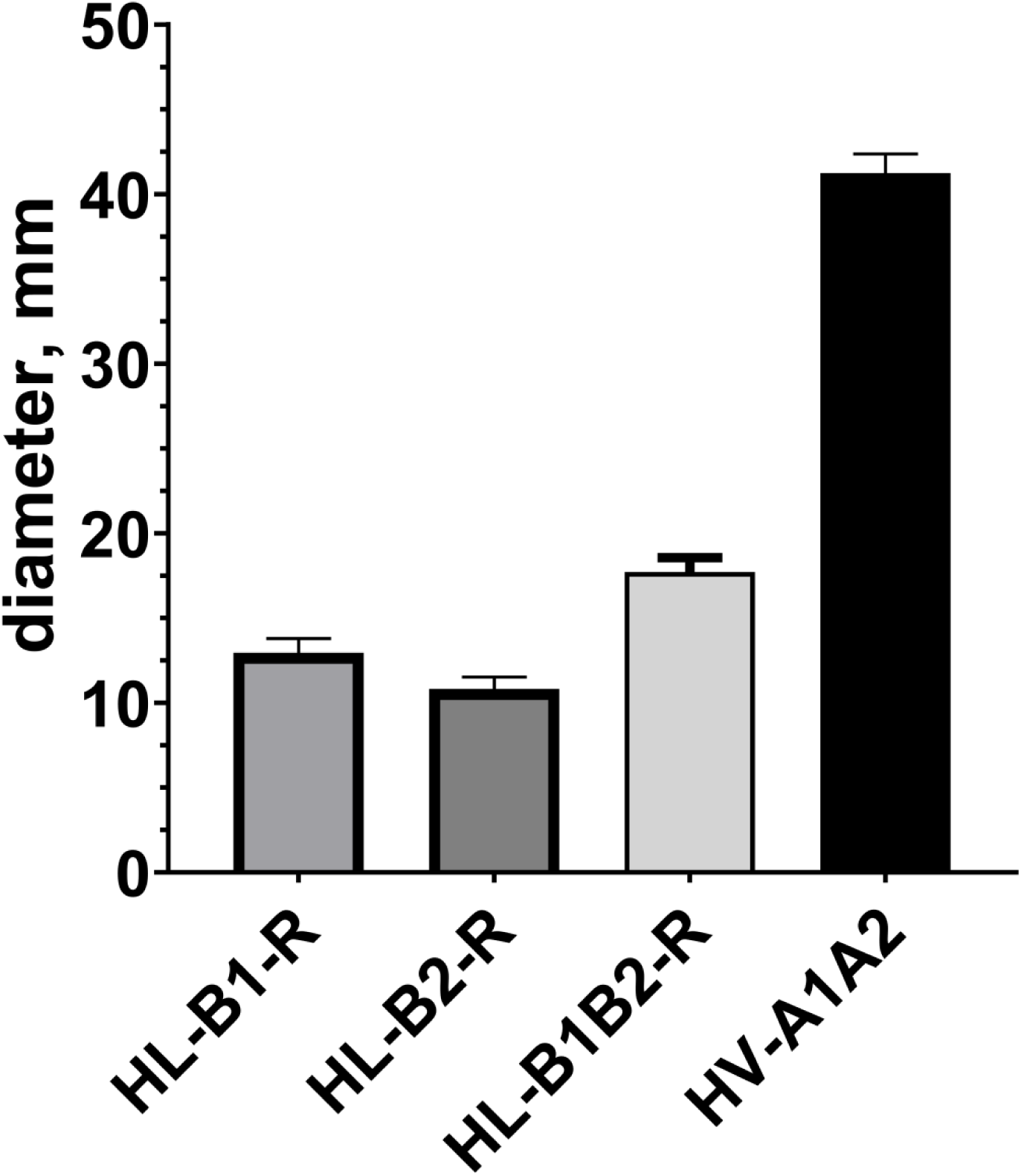
Comparison of cell motility of *Hfx. volcanii* strains expressing *Hrr. lacusprofundi* archaellin genes. The swarming diameters were measured 120 hours after inoculation: *Hfx. volcanii* MT2 transformed with pMT21 (HV-A1A2), pAS5 (HL-B1B2-R), pAS6 (HL-B1-R) and pAS7 (HL-B2-R), Mod-HV medium containing 0.5 mg/ml tryptophan, 0.24% agar.

**Figure 5.**
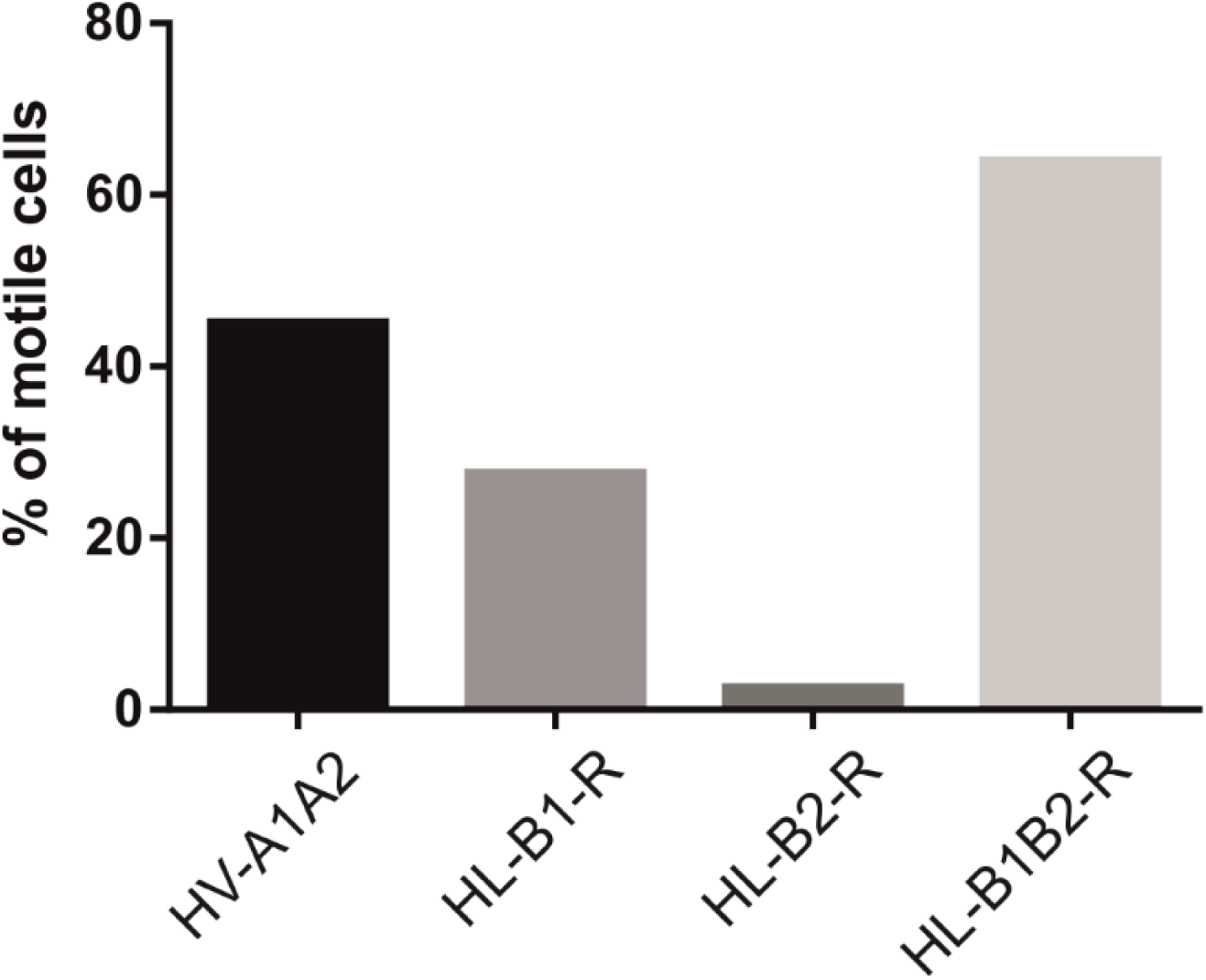
Swimming behavior of *Hfx. volcanii* strains expressing different combinations of archaellins. Percentage of motile cells in liquid cultures in early exponential phase. Cells were analyzed by light microscopy and time lapse images were taken for 15 seconds. All strains were analyzed at the same time during at least three independent experiments, which included >100 cells. The number of cells that showed motility during a 15 sec time lapse movie, was calculated as the percentage of the total observed cells. HL, archaellin from *Hrr. lacusprofundi.* HV, archaellin from *Hfx. volcanii*.

For isolation of archaella, we grew *Hfx. volcanii* cells in modified medium used by Guan et al. as described in the Material and Methods (Guan *et al.*, 2012). This modified medium in combination with moderately high growth temperature (37 °C) allowed us to obtain a high yield of recombinant archaella from *Hfx. volcanii*. The electrophoretic mobilities of recombinant archaellins differed a little from those of natural archaellins (Figure 2). The molar ratio of FlaB1/FlaB2 determined by scanning of the SDS-PAGE gels is approximately 1.20±0.04, which is higher than for natural archaella (0.90±0.04) (Figure 2). Both FlaB1 and FlaB2 sequences contain putative N-glycosylation sites, N-X-S(T) (**Figure S1**). Glycosylation of natural and recombinant archaellins was confirmed by staining with Schiff’s reagent. Note that in both native and recombinant filaments, several bands appear below the main FlaB1 band. These could represent different glycoforms, as they are differently stained by Schiff’s reagent. The lowest band may correspond to the non-glycosylated form (Figure 2). The structure of *Hfx. volcanii* archaellin oligosaccharides is well known (Tripepi *et al*., 2010; 2012). This is in contrast to the glycans of *Hrr. lacusprofundi*. In other *Halorubrum* species unique sialic acid-like saccharides not characteristic of the *Hfx. volcanii* were found (Zaretsky *et al*., 2018). Thus, despite the possible unnatural glycosylation in *Hfx. volcanii*, the *Hrr. lacusprofundi* archaellins retained their ability to assemble into functional archaella.

Comparison of electron microscopic images of native (HL-B2-N, HL-B1B2-N) and recombinant (HL-B2-R, HL-B1B2-R) archaella shows that recombinant archaella in general have the same features as natural ones (Figure 3). In electron micrographs, *Hrr. lacusprofundi* archaella appear as semi-flexible filaments in contrast to halobacterial archaella (Alam and Oesterhelt, 1984), which appear to have a more rigid helical shape. The HL-B1B2-R filaments have a wavy shape, characteristic for “classical” archaella (Figure 3). HL-B2-R archaella are also very flexible and often fold into loops, as is the case for the natural B2-N archaella. The HL-B1-R filaments appear straight and inflexible in comparison with the HL-B1B2-R archaella (Figure 3). This altered morphology of the HL-B1-R filaments might lead to motility problems. Indeed, life imaging showed that the percentage of motile cells expressing HL-B1-R is slightly reduced compared to HL-B1B2-R. The diameter of the recombinant and natural filaments is about the same (∼10 nm).

Since the *Hfx. volcanii* can exist in a fairly wide range of salinity (1.5 to 4 M NaCl) (Mullakhanbhai and Larsen, 1975), we decided to compare the functionality of various types of recombinant archaella depending on the content of sodium chloride in the growth medium. Motilities on semi-liquid agar were compared at four salinities: 10, 15, 20 and 25% NaCl (1.71 – 4.28 M) for *Hfx. volcanii* strains synthesizing the three archaella types (HL-B1-R, HL-B2-R, HL-B1B2-R). Interestingly, the HL-B1B2-R strain has an advantage in motility at 15 and especially at 10% NaCl (Figure 6). Thus, the two-component archaella appear to be better adapted to environmental salinity changes than the one-component archaella, allowing the species to occupy a the larger ecological niche.

**Figure 6.**
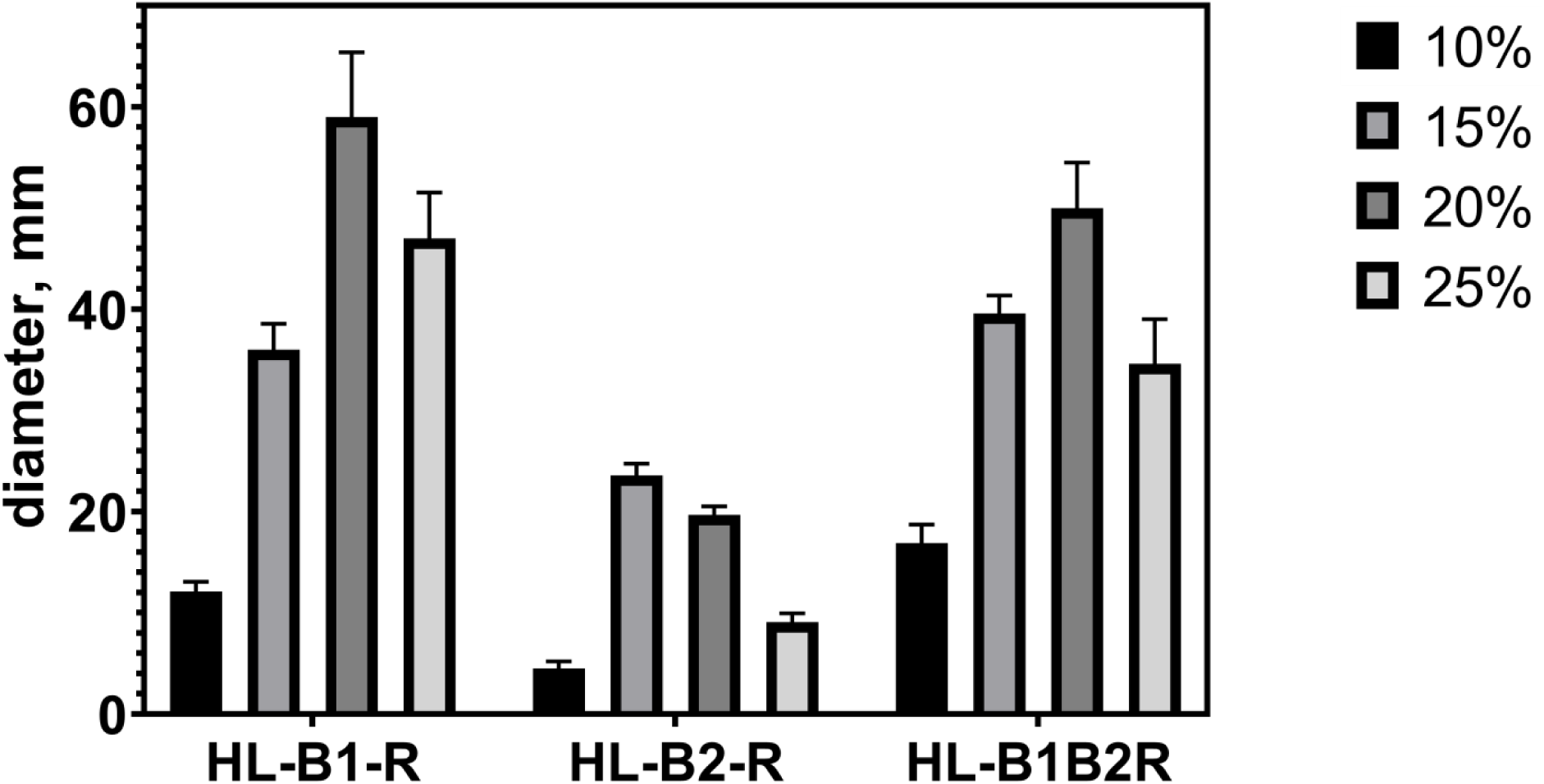
Comparison of cell motility of *Hfx. volcanii* strains expressing *Hrr. lacusprofundi* archaellins on semisolid media at different salinities. *Hfx. volcanii* MT2 strain was transformed with pAS5 (HL-B1B2-R), pAS6 (HL-B1-R) and pAS7 (HL-B2-R), Mod-HV medium containing 10, 15, 20 and 25% NaCl with 0.5 mg/ml tryptophan, 0.24% agar, 37 °C, 7 days.

*Halorubrum saccharovorum* ATCC 29252 (Tomlinson and Hochstein, 1976) is closely related to *Hrr. lacusprofundi* DL18 and has the same organization of the archaellin genes, encoding proteins with high similarity to those of DL18 (the identity of their FlaB1 archaellins is 70%, and FlaB2 - 73%). We isolated the archaella from *Hrr. saccharovorum* and established that, similar to *Hrr. lacusprofundi* DL18, in this organism both archaellins are present in an approximate 1:1 ratio also (**Figure S5**). We were able to successfully express the *Hrr. saccharovorum flaB1* gene and the *flaB1/flaB2* archaellin operon in *Hfx. volcanii* MT2. Unfortunately, an attempt to heterologously express the *Hrr. saccharovorum flaB2* gene did not lead to the archaella synthesis. An attempt was made to express the modified *Hrr. saccharovorum flaB2* gene in which the nucleotide sequence encoding the signal peptide was replaced with that for the FlaB1, but this also was unsuccessful. Possibly, FlaB2 of *Hrr. saccharovorum* can only form a stable filament in the presence of FlaB1. Probably, the *Hrr. saccharovorum* FlaB2 is less adaptable to heterogeneous assembly and a divergent glycosylation system. Synthesis of recombinant HS-B1-R and HS-B1B2-R archaella was confirmed by mass spectrometry (**Figure S15-S16**). The results of heterologous archaellin expression were similar to those found for recombinant expression of *Hrr. lacusprofundi* archaellins: expression of *flaB1/flaB2* and *flaB1* restored motility (**Figure S6**), and the synthesized archaella had a similar morphology to the native filaments (**Figure S8**). Again, the HS-B1-R filaments appeared straighter and less flexible as the HS-B1B2-R filaments (**S8**). Glycosylation of natural and recombinant archaellins of *Hrr. saccharovorum* was confirmed by specific staining with Schiff’s reagent (**Figure S5**). Interestingly, the staining intensities of natural *Hrr. saccharovorum* archaellins are noticeably less than that of recombinant archaellins. At the same time, both are apparently glycosylated less than natural *Hrr. lacusprofundi* archaellins.

### Scanning microcalorimetry experiments

To obtain additional information regarding archaella of different composition, we applied differential scanning microcalorimetry (DSC). Isolated archaella were heated in near natural (20% NaCl) and low (10% NaCl) salt conditions. We found that at 20% NaCl the two-component *Hrr. lacusprofundi* DL18 archaella (HL-B1B2-N) are much more stable than the natural HL-B2-N-archaella of *Hrr. lacusprofundi* ACAM 34. The temperature of the heat absorption peak maximum (T_m_) of the archaella consisting of two different archaellins (97.5 °C), was substantially higher than those of archaella build of a single type of archaellin (80.0 °C) (Figure 7). For both types of archaella only a single heat absorption peak was observed in the 20% NaCl buffer. This again indicates that the DL18 strain presents only a single type of archaella (consisting of both FlaB1 and FlaB2) at its surface. If two different types of filaments would be present, we would expect two different melting peaks. A decrease in the NaCl concentration (10%) resulted in a decrease of the T_m_. In 10% NaCl we observed a melting curve with two peaks (39 and 45 °C) for the one-component archaella HL-B2-N and single peak corresponding to a significantly higher temperature (81.5 °C) for the two-component archaella HL-B1B2-N (Table 3). These data suggest cooperative interactions and a close relationship of FlaB1 and FlaB2 subunits in the archaellar structure.

**Figure 7.**
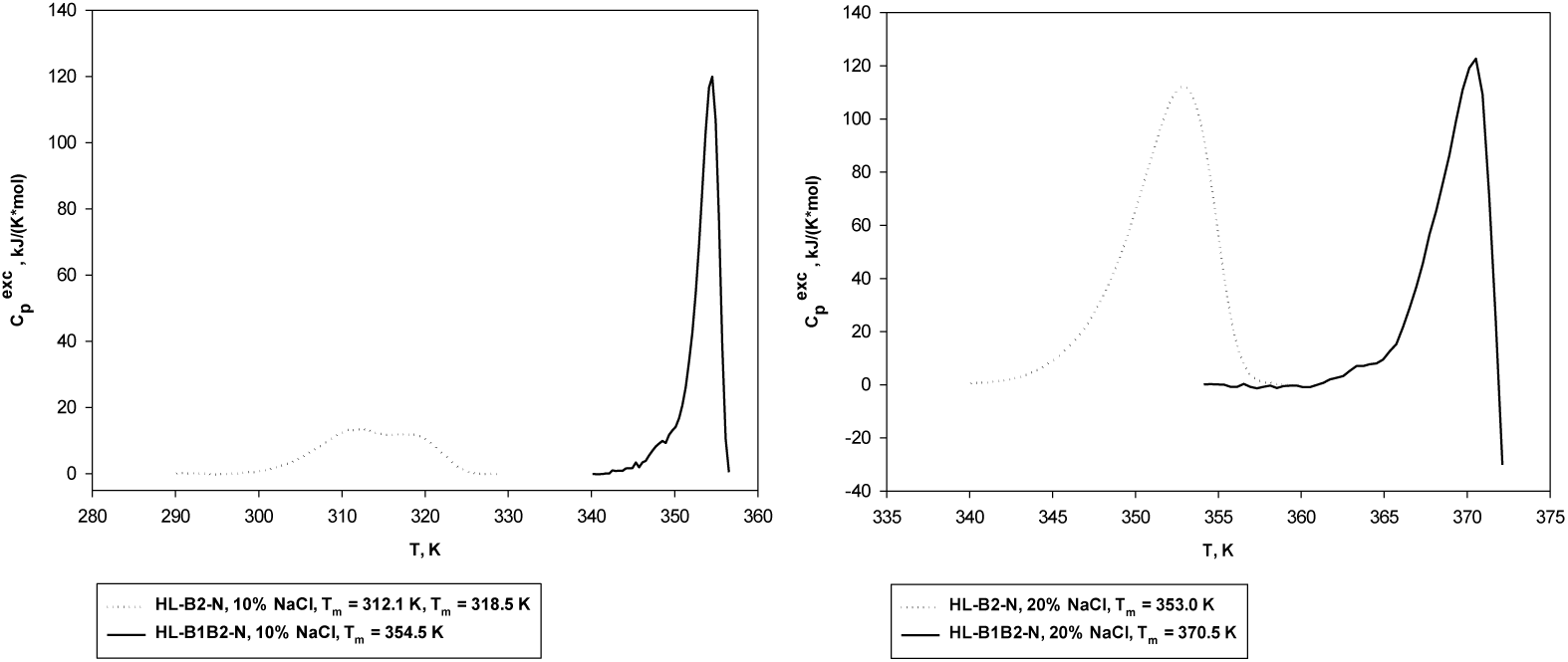
Temperature dependence of excess heat capacity of *Hrr. lacusprofundi* ACAM34 and DL18 archaellar filaments at two salinities: 20% (3.4 M) and 10% (1.7 M) NaCl, 10 mM Na-phosphate, pH 8.0.

**Table 1:** Plasmids and strains.

| Plasmid/Strain | Relevant properties | Reference |
| --- | --- | --- |
| Plasmids |  |  |
| pTA1228 | Amp<sup>r</sup>, *pyrE2* and *hdrB* markers, inducible *ptna* promoter | Brendel *et al.*, 2014 |
| pMT21 | pTA963 (Amp<sup>r</sup>, *pyrE2* and *hdrB* markers, inducible *ptna* promoter) containing *flgA1flgA2His* | Tripepi *et al.*, 2013 |
| pAS1 | pTA1228 containing *flaB1B2* of *Hrr. saccharovororum* | This study |
| pAS2 | pTA1228 containing *flaB1* of *Hrr. saccharovororum* | This study |
| pAS3 | pTA1228 containing *flaB2* of *Hrr. saccharovororum* | This study |
| pAS4 | pTA1228 containing *flaB2* of *Hrr. saccharovororum* (signal peptide was replaced to the FlaB1) | This study |
| pAS5 | pTA1228 containing *flaB1B2* of *Hrr. lacusprofundi* DL18 | This study |
| pAS6 | pTA1228 containing *flaB1* of *Hrr. lacusprofundi* DL18 | This study |
| pAS7 | pTA1228 containing *flaB2* of *Hrr. lacusprofundi* DL18 | This study |
| Strains |  |  |
| DH5α | *E. coli* F<sup>-</sup> ϕ80*lacZDM15* (*lacZYA-argF*) U169 *recA1 endA1 hsdR17 (rK<sup>-</sup> mK<sup>+</sup>) phoA supE44 thi-1 gyrA96 relA1* | Invitrogen |
| MT2 | *Hfx. volcanii* Δ*pyrE2* Δ*trp* Δ*flgA1*Δ*flgA2* | Tripepi *et al.*, 2010 |
| MT45 | MT2 containing pMT21 | Tripepi *et al.*, 2013 |
| AS1 | MT2 containing pAS1 | This study |
| AS2 | MT2 containing pAS2 | This study |
| AS3 | MT2 containing pAS3 | This study |
| AS4 | MT2 containing pAS4 | This study |
| AS5 | MT2 containing pAS5 | This study |
| AS6 | MT2 containing pAS6 | This study |
| AS7 | MT2 containing pAS7 | This study |
| ACAM 34 | *Halorubrum lacusprofundi* (B-1753, ATCC 49239) | Franzmann *et al.*, 1988 |
| DL18 | *Halorubrum lacusprofundi* | Erdmann *et al.*, 2017 |
| ATCC 29252 | *Halorubrum saccharovororum* (B-1747) | Tomlinson and Hochstein, 1976 |

**Table 2:**
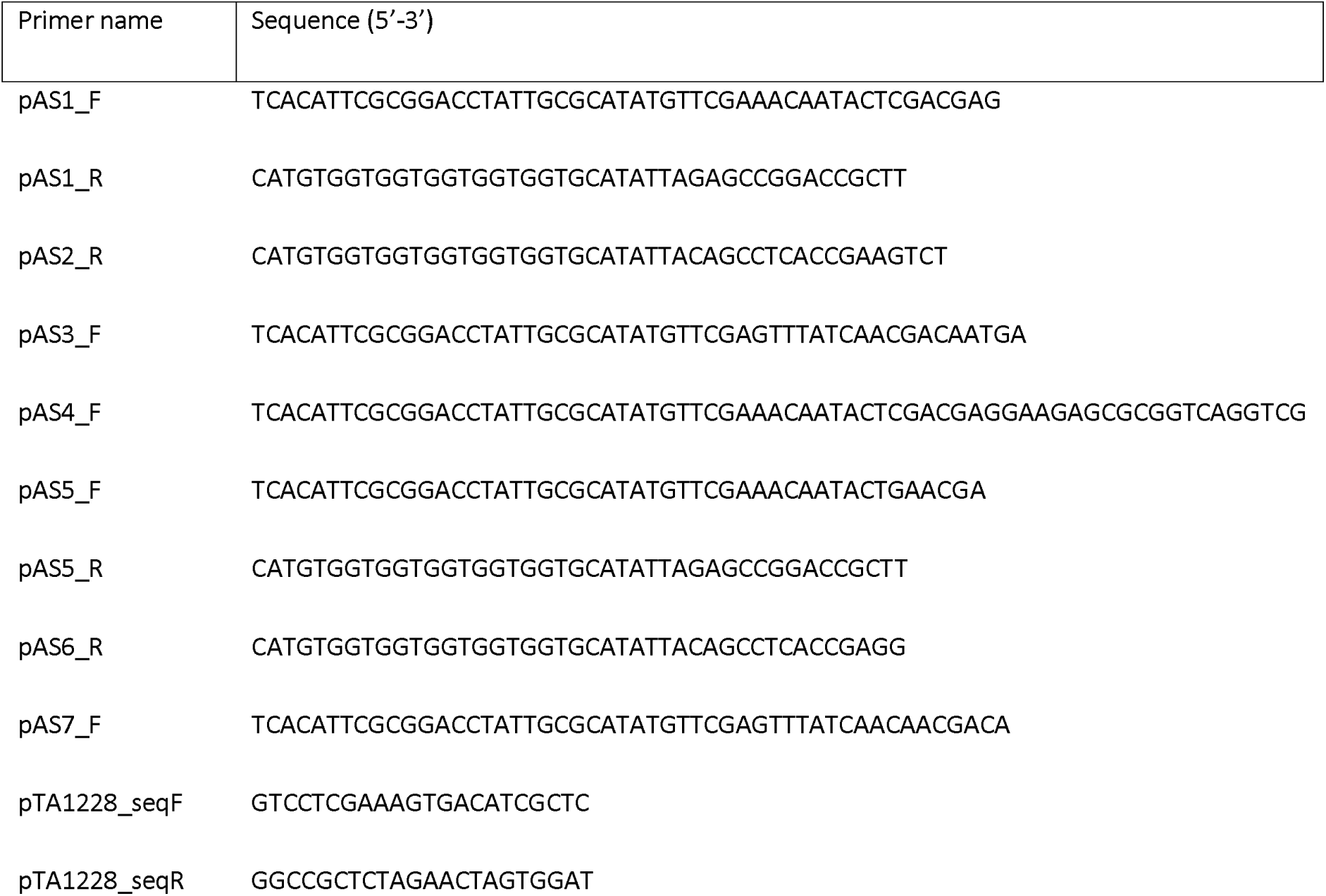
Primers.

**Table 3:** The melting points of different filament types, determined using scanning microcalorimetry.

| % NaCl | Archaeella | T<sub>m</sub> (°C) |
| --- | --- | --- |
|  | *Hrr. lacusprofundi* |  |
| 20 | B1B2-N | 97.5 |
| 20 | B2-N | 80.0 |
| 20 | B1B2-R | 92.3 |
| 20 | B1-R | 85.9 |
| 20 | B2-R | 74.3 |
| 10 | B1B2-N | 81.5 |
| 10 | B2-N | 39.1/45.5 |
| 10 | B1B2-R | 79.1 |
| 10 | B1-R | 71.4 |
| 10 | B2-R | 42.0 |
|  | *Hrr. saccharovororum* |  |
| 10 | B1B2-N | 85.2 |
| 10 | B1B2-R | 82.3 |
| 10 | B1-R | 81.9 |
| 5 | B1B2-N | 73.2 |
| 5 | B1B2-R | 71.9 |
| 5 | B1-R | 70.9 |

DSC data for recombinant filaments are similar to results obtained for the natural archaella. The temperature of the heat absorption peak maximum of HL-B1B2-R archaella (92 °C) under 20% NaCl was noticeably higher than that of recombinant filaments consisting either only of FlaB1 (86 °C) or FlaB2 (74 °C) (Figure 8). These temperatures were all slightly lower than those observed for the natural filaments isolated from *Hrr. lacusprofundi* (HL-B1B2-N, 97.5 °C and HL-B2-N, 80 °C). On melting at 10% NaCl extended heat absorption peak with a maximum of about 42 °C was observed for the HL-B2-R archaella (in comparison with two peaks at 39 and 45 °C for HL-B2-N). The HL-B1-R and HL-B2-R melting curves are different (Figure 8). Combination of both subunits in one archaellum filament (HL-B1B2) leads to structural changes that are reflected in a new melting curve, both natural and recombinant archaella.

**Figure 8.**
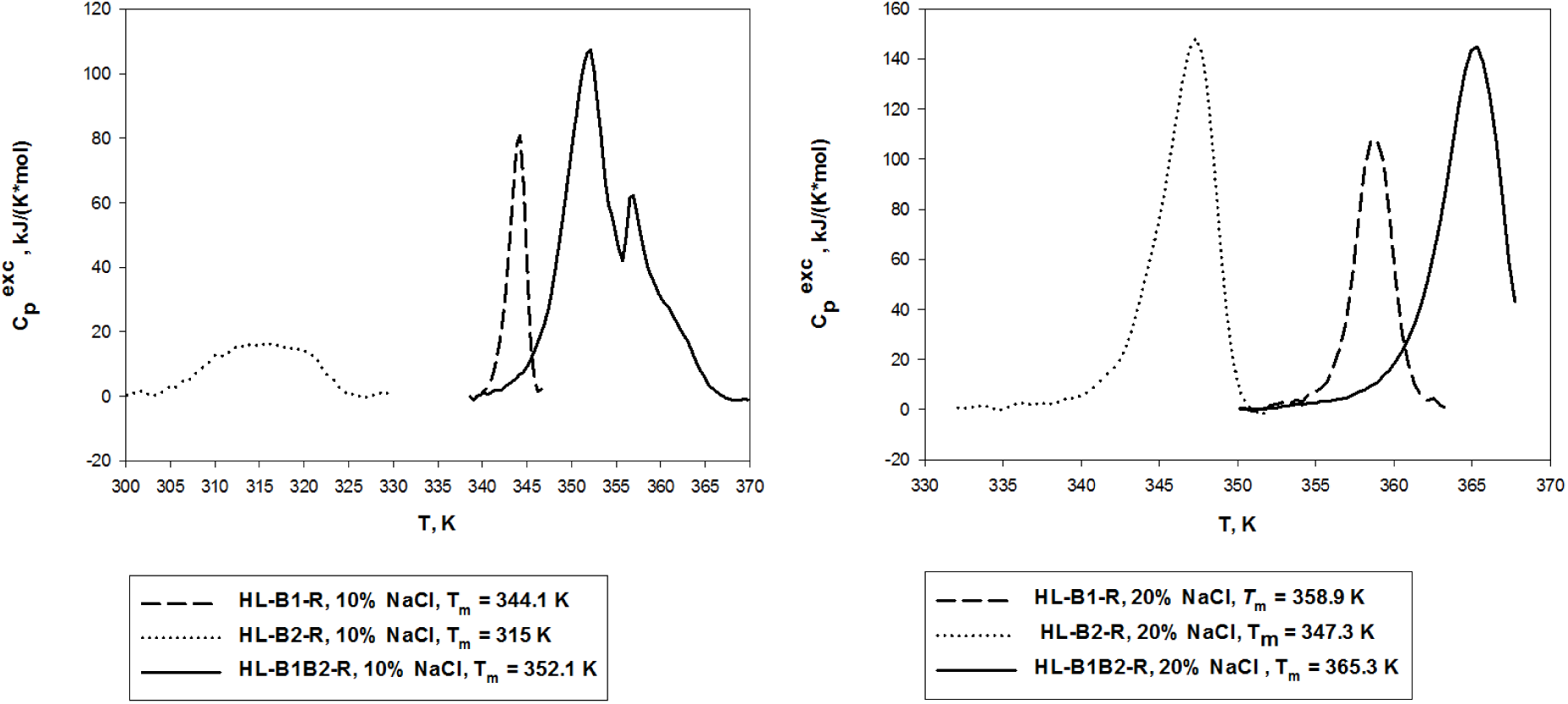
Temperature dependence of excess heat capacity of recombinant B1B2, B1 and B2 archaellar filaments at two salinities: 10% (1.7 M) and 20% (3.4 M) NaCl, 10 mM Na-phosphate, pH 8.0.

Similar experiments were carried out on *Hrr. saccharovorum* archaella. Here we have compared three archaella types: HS-B1B2-N, HS-B1B2-R and HS-B1-R. The experiments were carried out at 5 and 10% NaCl, since at 20% NaCl the natural *Hrr. saccharovorum* archaella melted near the upper limit of the experimental temperature range. In this case, the T_m_ of all three types of filaments were very similar, which is in line with the findings for *Hrr. lacusprofundi* archaella (**Figure S9**).

Thus, in general the B1B2 filaments are slightly more stable than the B1 and much more stable than the B2 filaments.

### Limited proteolysis confirms the interaction of *Hrr. lacusprofundi* archaellins in the filaments

To test the interaction between the FlaB1 and FlaB2 protein, *Hrr. lacusprofundi* archaellar filaments were subjected to trypsin digestion. FlaB2 from HL-B2-N archaella was cleaved by trypsin when the NaCl concentration was < 8%, while both FlaB1 and FlaB2 archaellins from HL-B1B2-N-filaments were protected from trypsin digestion under the same conditions (Figure 9). This effect was also observed for recombinant archaella. The HL-B1B2-R and HL-B1-R filaments are more resistant to trypsin digestion than the HL-B2-R (Figure 9). When the HL-B1-R and HL-B2-R archaellar filaments were mixed, the bands on the SDS-gel indicated digestion of FlaB2, suggesting that the stabilizing role of the intermolecular FlaB1/FlaB2 interactions only occurs when a mixed filament is built (Figure 9). Trypsin cleaves peptides on the C-terminal side of lysine and arginine residues. These residues are rare in *Hrr. lacusprofundi* archaellins (5 and 3 arginines, no lysines, respectively, in processed FlaB1 and FlaB2). From the distribution of arginines in the FlaB2 protein (**Figure S1**) (R61, R83, R230), it can be concluded that in the presence of FlaB1, the R61 and R83 sites of FlaB2 are protected from trypsin attack. From a comparison with the known archaella structures, it can be expected that R61 and R83 are localized between N-terminal α-helix and β-strand 1, and within in β- strand 2, respectively (Poweleit *et al*., 2016). The presence of FlaB1 seems to protect these sites, possibly by shielding them for trypsin.

**Figure 9.**
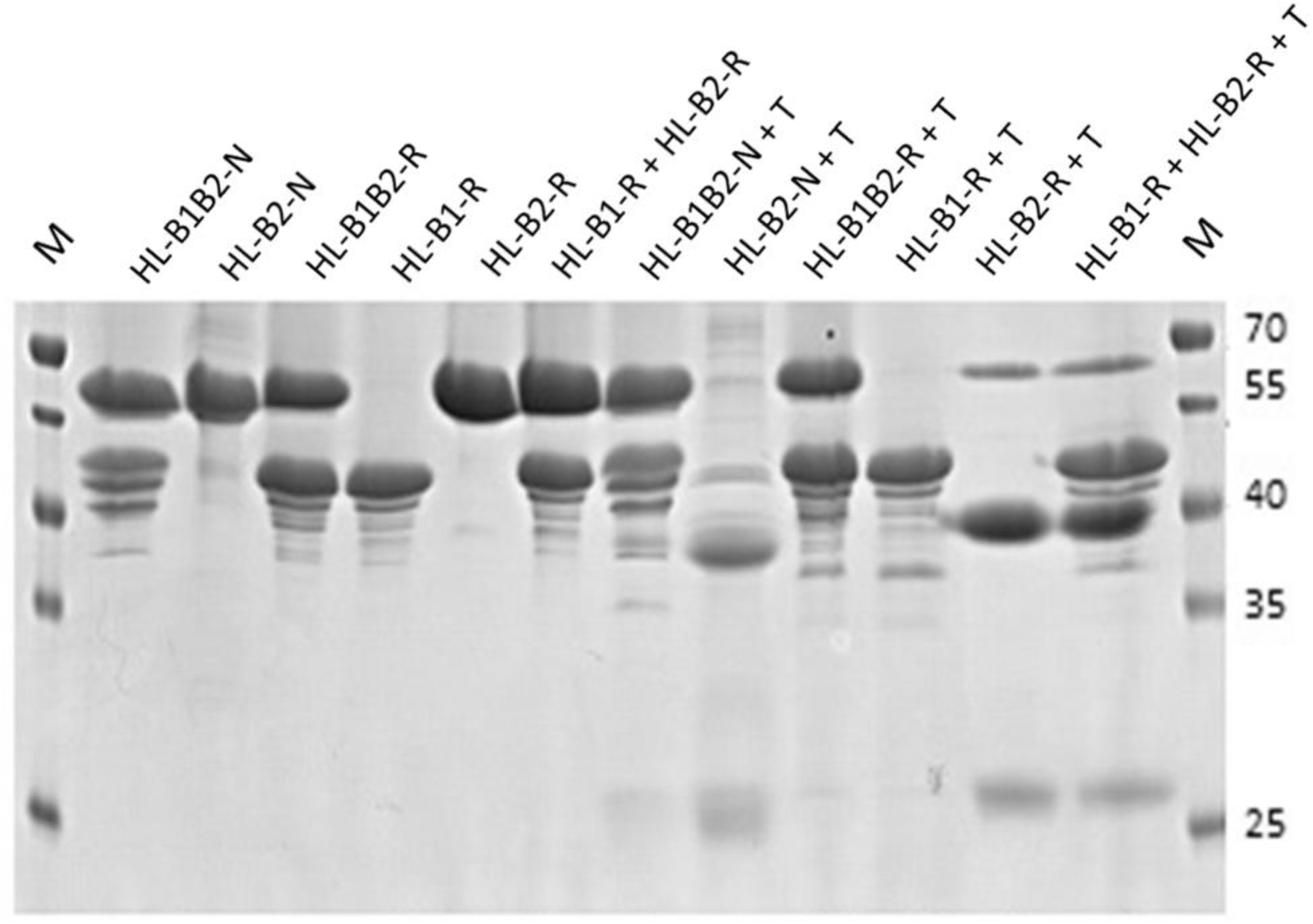
Results of limited trypsinolysis of isolated natural and recombinant *Hrr. lacusprofundi* archaellar filaments. Intact natural (HL-B1B2-N, HL-B2-N) and recombinant (HL-B1B2-R, HL-B1-R, HL-B2-R) archaella before and after trypsinolysis in 10 mM Na-phosphate, pH 8.0 containing 2% NaCl. Archaella were treated with trypsin 60 min at room temperature, the protein/enzyme ratio was 100:1. M - MW standards.

### Bioinformatical analysis of *Halorubrum* archaellins

After analyzing the *Halorubrum*s genomes available on insert date (50 in total), we found that in most species the organization of archaellin genes is similar to that of *Hrr. lacusprofundi* DL18: in 47 species there are two strongly diverged genes *flaB1* and *flaB2*, organized as a single operon, among them 9 species have an additional archaellin operon, which is likely to result from gene duplication events. Only *Hrr. lacusprofundi* ACAM 34 genome contains one archaellin gene. Each of the two *Halorubrum* archaellin types (FlaB1 and FlaB2) is characterized by a high degree of conservation. It is possible to identify generic signatures unique to each of two groups, for example, in internal (50-75 a. a.) and C-terminal partially conserved regions (**Figure S10**). FlaB1 sequences range from 187 to 210 amino acid residues, which is significantly shorter than the FlaB2 sizes from 201 to 456 a.a. Approximately 2/3^rds^ of the *Halorubrum* FlaB2 sequences have no more than 250 residues (**Table S1**). In most *Halorubrum* species the stop codon of gene *flaB1* and the start codon of *flaB2* are separated by a two nucleotide spacer CA (in 42 cases out of 53). The CC and CG spacers were found 3 times, AA, AC, AT and TT once

In two *Halorubrum* species (*Hrr. halodurans* and *Hrr. vacuolatum*) the organization of archaellin genes is fundamentally different: they contain several diverged archaellin genes that cannot be clearly classified as *flaB1* or *flaB2*. The corresponding proteins do not have signatures typical for archaellins of other *Halorubrum* species. Based on phylogenetic analysis, these archaellins should be probably attributed to the FlaB1 branch (Figure 10). The archaellin paralogs of these two species are more similar than the FlaB1 and FlaB2 paralogs in other *Halorubrum* species. Thus, identities between three *Hrr. halodurans* archaellins are > 55%, and >60% for four *Hrr. vacuolatum* archaellins. *Hrr. halodurans* archaellin genes constitute a single operon, the *flaB_a_* and *flaB_b_* genes are separated by CG spacer, the start codon of *flaB_c_* gene immediately follows the *flaB_b_* stop codon. The *Hrr. vacuolatum* genome contains three archaellin operons, one of them consists of two genes separated by a spacer of four nucleotides (GACC).

**Figure 10.**
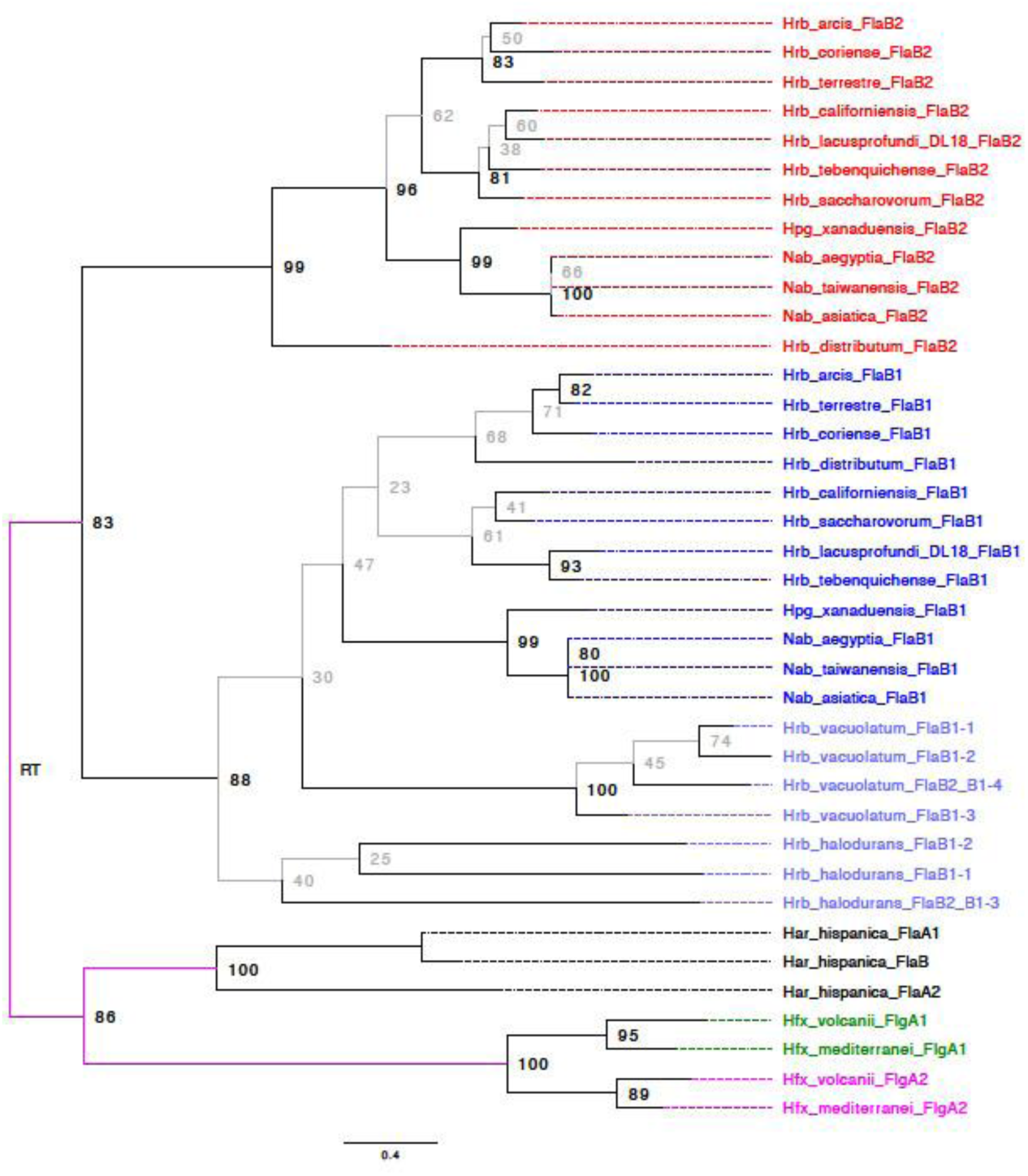
Phylogenetic tree of archaellins in the selected haloarchaea species (11 *Halorubrum* species, 2 species of *Halopiger* and *Natrialba*, *Hfx. volcanii* and *Har. hispanica*). Red – FlaB2 archaellins, Blue – FlaB1 archaellins. The depicted maximum likelihood phylogeny and bootstrap support values (traditional non-parametric) were calculated from a MAFT alignment without filtering for conserved sites using a WAG + F + G4 substitution model. Branches with less than 80% bootstrap support are given in grey. Branches from which the root cannot be excluded assuming that gene duplications occurred distal from the root are given in fuchsia. The depicted phylogeney was rooted using generalized midpoint optimization (Maljkovic Berry *et al.*, 2009).

Interestingly, for the other haloarchaeal genera (*Halopiger*, *Natrialba*, *Halobiforma*, *Natronolimnobius* and *Natrarchaeobius*) the situation with archaellins is very similar to that of *Halorubrum*. Despite the rather high similarity of their archaellins with that of the *Halorubrum* species, all these haloarchaea belong to a clade (the order *Natrialbales*, the family *Natrialbaceae*) evolutionarily distinct from *Halorubrum* (order *Haloferacales*, family *Halorubraceae*) (Amoozegar *et al.*, 2017). It is likely that operons from two highly diverged paralogs, having a common origin with *Halorubrum flaB1* and *flaB2* genes were exchanged between these taxa by horizontal gene transfer. The type of archaellins genes organization, characteristic for *Halorubrum*, predominates in genomes of the most representatives of these groups. It is interesting that, unlike *Halorubrum*, these haloarcheal groups are characterized by large FlaB1 archaellins (from 193 to 481 a.a., for half of them the archaellin size is >300 a.a.) and relatively small FlaB2 (from 208 to 261 a.a.). In genomes of some species (*Hpg. djelfimassiliensis* and *Hbf. nitratireducens*) archaellin genes have a type of organization, similar to that in *Hrr. halodurans* and *Hrr. vacuolatum*. The corresponding protein products have their own characteristics, which are not typical of most *Halorubrum* archaellins.

*Halorubrum* archaellins reveal similarity with archaellins of the evolutionarily distant (Becker *et al.*, 2014) haloarchaeal groups *Halobiforma*, *Halopiger*, *Natrialba* and *Natronolimnobius*. The evolutionary history of the archaellins clearly does not reflect organismal relationships inferred from genome comparisons (Gupta et al., 2016). *Halorubrum* is considered a member of the *Haloferacales*, whereas *Halobiforma*, *Halopiger*, *Natrialba* and *Natronolimnobius* are placed in the *Natrialbales*. Figures 10 **and S11** shows that for both *flaB1* and *flaB2* these divergent microorganisms form well supported clades in the archaellin phylogeny. Of particular interest is the recently described haloarchaea *Natrarchaeobius chitinivorans* (Sorokin *et al.*, 2019) having two archaellin operons. One operon consists of two genes closely related to *Hrr. lacusprofundi flaB1* and *flaB2*, and the other distant operon consists of three genes having a different origin.

It should be emphasized that, as can be seen from the evolutionary tree (Figure 10), the divergence between *flaB1* and *flaB2* genes is a more ancient compared to the divergence of the corresponding genes in such haloarchaea as *Hrr. halodurans*, *Hrr. vacuolatum*, *Hfx. volcanii*, *Har. hispanica* and *Hbt. salinarum*. *Hfx. volcanii* and *Hrr. lacusprofundi* DL18 represent two distinct types: the first of them (*Hfx. volcanii*) is characterized by archaellins that have a close relationship, and the second group is characterized by highly divergent archaellins. The occurrence of the second type among divergent groups of *Halobacteria* is likely due to horizontal gene transfer between *Natrialbales* and *Haloferacales*. It can be assumed that the principles of the structural organization of the archaella has significant differences between these two types.

## DISCUSSION

The archaeal motility structure, the archaellum, consists of thousands of copies of N-terminally cleaved archaellin subunits. While crenarchaea usually encode a single type of archaellin, the euryarchaea are characterized by the presence of multiple types of archaellin encoding genes. Recently, high resolution structures of archaellar filaments of methanogens and hyperthermophilic euryarchaea became available (Poweleit *et al.*, 2016; Daum *et al.*, 2017; Meshcheryakov *et al.*, 2019). In these structures only a single type of archaellin is present in the filament, even though several archaellin genes are present in the genome. Daum *et al*. suggested that these other archaellins either (i) are minor, and form specific basal or terminal segments of the filament, or (ii) that each of the different types of archaellins forms individual filaments (Daum *et al.*, 2017).

We aimed to understand the biological relevance of archaellin multiplicity by using the halophilic euryarchaeon *Hrr. lacusprofundi*, which encodes two divergent archaellins, FlaB1 and FlaB2, that are easily distinguishable from each other in amino acid sequences and sizes. Both archaellins were present in the archaellum filaments in comparable amounts. We used natural *Hrr. lacusprofundi* strains encoding FlaB1 and FlaB2 or only the FlaB2 protein. In addition, we expressed the FlaB1, FlaB2 and a combination of the two proteins in *Hfx. volcanii*. This allowed us to study the role of the individual archaellins, indicating that a combination of FlaB1 and FlaB2 is required to provide stability to the archaellum. In *flaB1* and *flaB2* containing strains, microcalorimetry and SDS-PAGE analysis confirmed that both archaellins are part of each filament. Differential melting curves in scanning microcalorimetry experiments and protection against trypsin digestion indicate that the FlaB1 and FlaB2 proteins tightly interact within the archaellum and as such provide stability to the filament. In the absence of one of the archaellins, the archaella either become over flexible (FlaB2 filaments), or quite stiff (FlaB1 filaments). These archaella consisting of single components are still functional and provide motility, although that of filaments consisting only of FlaB2 is strongly reduced. Comparison of the motility of *Hfx. volcanii* strains expressing different archaella types indicates that two-component archaella are better adapted to stress caused by both extra low and extra high salt concentrations.

We suppose that FlaB1 forms a filament “framework” and FlaB2 adopts a final more stable conformational state by interacting with FlaB1. It can also be assumed that FlaB1 and FlaB2 form a heterodimer that then assembles into the archaella. The pairwise interaction between FlaB1 and FlaB2 is compatible with the single higher melting point. The microcalorimetry experiments show that the archaella consisting of two types of archaellins are more stable under varying conditions, such as low salt stress. In a comparative study of one- and two-component filaments, we have found that the presence of FlaB1 substantially stabilizes the filament structure. Instead of a superposition of FlaB1 and FlaB2 peaks, we observe a new peak of heat absorption, that melting point was slightly higher than for FlaB1-filaments, and significantly higher than for FlaB2-filaments.

Thus, the two-component composition of *Hrr. lacusprofundi* archaellar filaments is needed for additional stabilization of the archaellum structure and adaptation to a wider range of external conditions and is not required for archaella supercoiling.

By applying heterologous expression of the *Hrr. lacusprofundi* archaellins in *Hfx. volcanii*, we demonstrated that archaellins can assemble in functional archaella, even in species that possess highly divergent archaellin genes. This suggests that foreign archaellin genes captured via horizontal transfer can quite easily adapt to the assembly and glycosylation system of the new host. Exchange of archaellins could provide an evolutionary advantage as it might allow adaptation to new environments or block the attachment of archaellum specific viruses (Pyatibratov *et al.*, 2008; Tschitschko *et al.*, 2018).

Euryarchaea are characterized by the genomic presence of multiple different archaellin genes. Most euryarchaea have two archaellin genes, but in some cases the number of different archaellins is even higher (such as *Hht. litchfieldiae*, which has seven archaellin genes) (Tschitschko *et al.*, 2015).

Several explanations for the existence of multiple archaellins have been proposed. Firstly, some archaellins might form minor components of the archaellum. This hypothesis is consistent with the results of the work (Chaban *et al.*, 2007), when the archaellum hook segment is built of the minor FlaB3 archaellin in the methanogenic archaea. Also in *Hfx. volcanii* functional archaella can be formed only from the major archaellin FlgA1, and cells with such archaella are hypermotile compared to cells of the natural strain, whose archaella consist of two archaellins (Tripepi *et al.*, 2013). Secondly, multiple archaellins were shown to act as ecoparalogs. *Har. marismortui* is capable of assembling functional (i.e., spiral) filaments from either one of the two encoded archaellins (Syutkin *et al.*, 2014). The type of archaellins incorporated in the filament is dependent on environmental conditions (such as ionic strength) (Syutkin *et al.*, 2019).

Archaella of several euryarchaea consist of multiple archaellins in comparable amounts, which is not in correspondence with either of the two above mentioned explanations. For example, the archaellar filaments of *Hbt. salinarum* (formerly *Hbt. halobium*) contain the products of all five archaellin genes, and the proportion of archaellins FlgA1 and FlgB2 is comparable with the proportion of archaellins FlgA2, FlgB1 and FlgB3 (Gerl *et al*., 1989). Earlier it had been suggested that archaellin multiplicity may cause the archaellum to became supercoiled (Tarasov *et al.*, 2000). This hypothesis was based on an analogy with the bacterial flagellar filaments, where two conformational flagellin states provide filament spiralization (Calladine, 1978). It was shown that inactivation of the *Hbt. salinarum* archaellin genes led to disruption of the archaella assembly, while only straight filaments could be formed from the product of a single archaellin gene (*flgA1* or *flgA2*) (Tarasov *et al.*, 2000; 2004). In case of *Hrr. lacusprofundi*, the presence of two archaellins is not required for spiralization, as functional filaments can be formed from each of the single archaellin types. However, the presence of two archaellins provides extra stability to the filament, causing it to better withstand ionic stress conditions, and to provide the highest level of motility.

Thus, with this work we add another aspect of encoding multiple archaellins to the other previously discovered mechanisms. Besides forming specialized minor components of the filament, or acting as ecoparalogs, we now show that multiple archaellins can also be important to form the filament with optimal properties in terms of flexibility and stability. In addition, we provide evidence that the exchange of archaellins between different species, can result in functional archaellum filaments. Together, these findings sketch an evolutionary picture in which accidental duplication of archaellins, was used to the advantage of several euryarchaea, specifically the haloarchaea. Frequent horizontal gene transfer of archaellins promotes the evolutionary adaptability of different species. In light of these evolutionary advantages, one might wonder how crenarchaea can be successful with only a single type of archaellin. Possibly the variation of environmental conditions of haloarchaea (such as ionic strength), promote the presence of multiple archaellins.

## EXPERIMENTAL PROCEDURES

### Strains and media

The plasmids and strains used in this study are listed in Table 1. The strains *Halorubrum lacusprofundi* B-1753 (ACAM 34, ATCC 49239) and *Halorubrum saccharovorum* B-1747 (ATCC 29252) from the All-Russian Collection of Microorganisms (VKM), Pushchino; strain *Halorubrum lacusprofundi* DL18 (Erdmann *et al.*, 2017) was kindly provided by R. Cavicchioli; strain *Haloferax volcanii* MT2 (Tripepi *et al.*, 2010) was kindly provided by M. Pohlschröder.

The *Hrr. lacusprofundi* cells were grown under moderate aeration at 37 °C in a liquid medium containing 0.5% casamino acids, 0.5% yeast extractcasamino acids, 3.42 M (20%) NaCl, 27 mM KCl, 80 mM MgSO_4_, 12 mM sodium citrate, 6 mM sodium glutamate, pH 7.2. Filter sterilized aqueous solution of microelements (1.7 mL) containing 0.9 mM MnCl_2_ · 7H_2_O and 17 mM FeCl_3_ · 7H_2_O was added to 1 L of the medium after autoclaving. All *Hfx. volcanii* transformed strains were grown at 37 °C in liquid or solid/semi-solid agar medium containing 0.5% casamino acids, 2.91 M (17%) NaCl, 0.15 M MgSO_4_, 1 mM MnCl_2_, 50 mM KCl, 3 mM CaCl_2_, pH 7.2 (Mod-HV medium) and 1.2 % (solid) or 0.24% (semi-solid) agar. Heterologous archaella synthesis was induced by addition of tryptophan (concentration of 0.2 - 1 mg/ml).

In experiments testing motility comparing at various salinity, we used Mod-HV media with NaCl concentrations of 10, 15, 20, and 25% (1.71, 2.57, 3.42 and 4.28 M), while the concentrations of the remaining components did not change. The swarming diameters on semi-solid agar plates were measured daily. Ten biological replicates were performed and average diameter values and standard deviations were calculated. Data were analyzed using GraphPad Prism 8.0 (GraphPad Software Inc., San Diego, USA).

To isolate recombinant archaella from liquid medium (with the same composition, but 0.5% casamino acids were replaced by 1% yeast extract) a piece of agar from the edge of the spot was used for inoculation.

Life cell imaging was performed in CA medium as described by Quax et al (Quax *et al.*, 2018). In short, cells were grown until an OD of ∼0.05 and imaged in a round DF 0.17 216mm microscopy dish (Bioptechs) and observed at 20x magnification in the PH2 mode with a Zeiss Axio Observer 2.1 Microscope equipped with a heated XL-5 2000 Incubator running VisiView® software.

Time lapse movies were recorded for 15 sec and the X,Y co-ordinates of cells were tracked with Metamorph® software. From the X,Y co-ordinates the average velocity of a given time frame was calculated for each cell using the Pythagoras Theorem. In addition the X, Y coordinates were used to calculate the number of turns larger than 90 ° that each cell made per second in the total time it was tracked. In case no 90° turn was observed within the time frame of the 15 sec movie, the cell was automatically assigned the value >16. Velocity and turns per second values were averaged for technical replicates recorded in a single experiment on a single day. Each experiment was performed at least on three independent occasions. The percentage of motile cells was calculated by counting cells that displayed motility in a 15 sec time lapse movie, and dividing this by the total number of visible cells in the frame of this movie. This was done for at least 10 movies (displaying ∼50 cells each) for each strain for each experiment. The experiment was performed on at least three independent occasions.

### Preparation of DNA and polymerase chain reaction (PCR)

The plasmids for heterologous archaellin expression were assembled by the SLIC method (Li and Elledge, 2012) with modifications. These expression vectors included the inducible tryptophanase promoter (tna) to drive expression of these genes. The DNA fragments, containing desired archaellin genes were amplified from *Hrr. lacusprofundi* or *Hrr. saccarovorum* genome with the primers described in Table 1. The PCR was performed with Q5 High-Fidelity DNA Polymerase (New England Biolabs), temperatures were as follows: 98 °C for 30 sec for initial denaturation, 25 cycles: 10 sec at 98 °C, 20 sec at 65 °C, 90 sec at 72 °C, and then 2 min at 72 °C for final elongation. Resulting PCR products were purified from reaction mixture and mixed with the pTA1228 vector preliminary linearized by NdeI treatment. Each assembly mix contained 100 fmol of both vector and insert, as well as 1.5 U of T4 DNA polymerase and NEBuffer 2.1 (New England Biolabs). The reaction was incubated for 4 minutes at 22 °C and then stopped by addition of 10 mM dCTP. The resulting mix was used for *E. coli* transformation. The colonies appeared at the next day were analyzed by PCR with the pTA1228_seqF and pTA1228_seqR primers. The plasmids from positive colonies were isolated and correct assembly of the plasmid was confirmed by sequencing with the primers used for colonies screening.

### Isolation of archaellar filaments

Archaellar filaments were prepared by precipitation with polyethylene glycol 6000 (Gerl *et al.*, 1989). The protein preparations were dissolved in 10 mM Na-phosphate, pH 8.0 containing appropriate NaCl concentrations (0-20%) at a concentration of 0.5-1.0 mg/ml. SDS-PAGE was performed using 9-12% acrylamide gels. The proteins were stained with Coomassie brilliant blue G 250. To prepare samples for microcalorimetry, the archaellar filaments were also dialyzed against the above-mentioned buffer solutions. Protein concentrations were determined using the Coomassie Plus Protein Assay Reagent kit (Pierce, IL) according to the manufacturer’s protocol. ImageJ software (NIH) was used to scan stained acrylamide gels and estimate FlaB1/FlaB2 ratios.

### Chromatography mass spectrometry analysis

Proteins bands were excised and treated with Proteinase K (Promega) and trypsin (Sigma) at 37°C in a Thermo Mixer thermo shaker (Eppendorf, Germany). To stabilize proteinase K, CaCl_2_ was added to the solution to a final concentration of 5 mM. The molar ratio of enzyme-to-protein was 1/50. The reaction was stopped by adding trifluoroacetic acid to the solution. Prior to mass spectrometric analysis, the peptides were separated by reversed-phase high-performance liquid chromatography using an Easy-nLC 1000 Nanoliquid chromatography (ThermoFisher Scientific). The separation was carried out in a homemade column 25 cm in length and 100 μm in diameter packed with a C18 adsorbent; with an adsorbent particle size of 3 μm and a pore size of 300 Å. The column was packed under laboratory conditions at a pressure of 500 atm. The peptides were eluted in a gradient of acetonitrile from 3% to 40% for 180 min; the mobile phase flow rate was 0.3 μl/min. Mass spectra of the samples were obtained using an OrbiTrap Elite mass spectrometer (Thermo Scientific, Germany). The peptides were ionized by electrospray at nano-liter flow rates with 2 kV ion spray voltage; ion fragmentation was induced by collisions with an inert gas (collision induced dissociation in a high-energy cell). The mass spectra were processed and peptides were identified using Thermo Xcalibur Qual Browser and PEAKS Studio (ver. 7.5) programs based on the sequences of UniRef-100. Parent Mass Error Tolerance was 2.0 ppm and fragment Mass Error Tolerance was 0.1 Da. Only peptides were taken into account with a “10 l gP.” threshold value higher than 15.

### Electron microscopy

The archaellar filament specimens were prepared by negative staining with 2% uranyl acetate on Formvar-coated Copper grids. A grid was floated on a 20-μl drop of filament solution (about 0.01 mg/ml, in 20% NaCl, 10 mM Na-phosphate, pH 8.0) for 2 min, blotted with filter paper, placed on top of a drop of 2% uranyl acetate and left for 1-1.5 min. Excess stain was removed by touching the grid to filter paper, and the grid was air dried. Samples were examined on a Jeol JEM-1400 transmission electron microscope (JEOL, Japan) operated at 120 kV. Images were recorded digitally using a high-resolution water-cooled bottom-mounted CCD camera.

### Scanning microcalorimetry

Scanning microcalorimetry experements were performed on a differential scanning microcalorimeter SCAL-1 (Scal Co., Pushchino, Russia) with a 0.33 glass cell at a heating rate of 1 K/min, under a pressure of 2.5 atm (Senin et al., 2000). The measurements and necessary calculations were performed according to (Privalov and Potekhin, 1986) and described in detail in (Tarasov *et al.*, 1995).

### Limited proteolysis

Limited proteolysis by trypsin (Sigma) was performed in 10 mM Na-phosphate buffer, pH 8.0 at 21 °C. 20 μl aliquots for electrophoresis were taken at defined time periods. The reaction was terminated by adding an equimolar amount of trypsin inhibitor from ovomucoid (Sigma).

### Phylogenetic reconstruction

Sequences were aligned with MAFFT (Katoh et al., 2013) or PRANK (Löytynoja et al., 2008). For some analyses unreliably aligned sites were removed using guidance (Sela et al., 2015). Search for the best model to describe sequence evolution and search for the maximum likelihood tree were performed in IQ-TREE (Nguyen et al., 2015) using the Bayesian Information Criterion (BIC). The only difference between phylogenetic reconstruction from the different alignments is that in case of alignments filtered for conserved sites, the branches are shorter (due to the removal of variable sites), the bootstrap support values are lower, and the most appropriate models determined with IQ-TREE are simpler, because the removal of the variable more difficult to align sites also removes phylogenetic information.

## Acknowledgements

This work was supported by the RFBR grant No. 19-04-00613 A to M.P and an Emmy Noether grant (411069969) from the Deutsche Forschungs Gemeinschaft to TEFQ. We thank Dr. T Allers for pTA1228 plasmid, Dr. R. Cavicchioli and Dr. M. Pohlschröder for the kindly presented strains and M. Suvorina for the help with MS analysis. Electron microscopy was performed with the support of Moscow State University development program (PNR 5.13). Mass spectrometric analysis was performed in facilities of United Pushchino Center “Structural and functional studies of proteins and RNA” (584307).

## Author contributions

M.G.P, A.S.S., T.E.F.Q., T.N.M. and J.P.G. conceived of the project, designed the study, and wrote the paper; M.G.P, A.S.S., T.E.F.Q., T.N.M., I.I.K., and A.K.S. performed the experiments; M.G.P, A.S.S., T.E.F.Q., T.N.M., R.T.P., J.P.G. and O.V.F. analyzed the data; S.N.B. and A.V.G. contributed to experiments; M.G.P, A.S.S., T.E.F.Q., I.I.K., A.K.S. and O.V.F. contributed funding and resources.

## SUPPLEMENTARY FIGURE LEGENDS

**Figure S1.** Alignment of the sequence of the ACAM 34 and DL18 *Hrr. lacusprofundi* archaellins. Sequence alignment was performed using Clustal Omega (Sievers and Higgins, 2018; http://www.clustal.org/omega/). In each sequence, potentially N-glycosylation sites, N-X-T(S), are boxed. The signal peptide cleavage site is indicated by an arrow.

**Figure S2.** Nucleotide sequences of archaellin genes of *Hrr. lacusprofundi* DL18 and ACAM 34 strains. The sequence of the *flaB1* gene is highlighted in yellow, and *flaB2* gene 2 - in green. The sequences that are only in *flaB2* of the DL18 and are absent in that of ACAM 34 are highlighted in red. Sequences common to both *flaB1* and *flaB2* of the ACAM 34 are highlighted in blue.

**Figure S2.** Nucleotide sequences of archaellin genes of *Hrr. lacusprofundi* DL18 and ACAM 34 (ATCC 49239) strains. The sequence of the *flaB1* gene is highlighted in yellow, and *flaB2* gene 2 - in green. The sequences that are only in *flaB2* of the DL18 and are absent in that of ACAM 34 are highlighted in red. Sequences common to both *flaB1* and *flaB2* of the ACAM 34 are highlighted in blue.

**Figure S3.** Swimming behavior of *Hfx. volcanii* strains expressing different combinations of archaellins. Liquid cultures were analyzed by light microscopy at 45 °C. (*A*) Average velocity of different strains. Bars indicate SD. (*B*) Tukey box plot of number of seconds between two subsequent turns of >90°. For each strain, the middle line in the box displays the median. Boxes display the 25–75th percentile, and lower and upper bars represent the minimum and maximum time. HL, archaellin from *Hrr. lacusprofundi.* HV, archaellin from *Hfx. volcanii*.

**Figure S4.** Electron micrographs of *Hfx. volcanii* MT2 transformed with pAS5 (HL-B1B2-R) (top), pAS6 (HL-B1-R) (middle) and pAS7 (HL-B2-R) (bottom) negatively stained with 2% uranyl acetate. Scale bars = 1 μm.

**Figure S5.** Archaellin staining with Schiff’s reagent in 12.5% polyacrylamide gel (right); the same gel stained with Coomassie G250 (left). M - prestained protein standard, HL-B1B2-N and HL-B1B2-R – natural and recombinant *Hrr. lacusprofundi* DL18 archaella, HS-B1B2-N – natural archaella of *Hrr. saccharovorum*, HS-B1B2-R, HS-B1-R – recombinant *Hrr. saccharovorum* archaella isolated from *Hfx. volcanii* MT2: A1A2 – *Hfx. volcanii* MT45 archaella (positive control), BSA - bovine serum albumin (negative control).

**Figure S6.** Comparison of cell motility of *Hfx. volcanii* strains expressing *Hrr. lacusprofundi* and *Hrr. saccharovorum* archaellin genes. Photos of swarming plates were taking at different times after inoculation. Top: left – *Hfx. volcanii* MT2 transformed with pMT21 (HV-A1A2), pAS6 (HL-B1-R), pAS7 (HL-B2-R) and pAS5 (HL-B1B2-R), Mod-HV medium containing 0.5 mg/ml tryptophan, 0.24% agar, 37 °C, 5 days; right – the same, without pMT21, 8 days. Bottom: left – *Hfx. volcanii* MT2 transformed with pMT21 (FlgA1FlgA2), pAS2 (HS-B1-R), pAS3 (HS-B2-R) and pAS1 (HS-B1B2-R), the above-mentioned medium, 5 days; right – the same, without pMT21, 8 days.

**Figure S7.** Comparison of cell motility of haloarchaeal strains expressing *Hrr. saccharovorum* archaellins. The swarming diameters were measured 120 hours after inoculation: *Hfx. volcanii* MT2 transformed with pMT21 (HV-A1A2), pAS1 (HS-B1B2-R), pAS2 (HS-B1-R) and pAS3 (HS-B2-R), Mod-HV medium containing 0.5 mg/ml tryptophan, 0.24% agar.

**Figure S8.** Negatively stained (1 % uranyl acetate) preparations of natural (HS-B1B2-N) and recombinant (HS-B1B2-R and HS-B1-R) archaellar filaments of *Hrr. saccharovorum* in 20% NaCl, 10 mM Na-phosphate, pH 8.0. Scale bar – 100 nm.

**Figure S9.** Temperature dependence of excess heat capacity of *Hrr. saccharovorum* recombinant HS-B1B2-R and HS-B1-R archaellar filaments in comparison with native (HS-B1B2-N) filaments at two salinities: 10% (1.7 M) and 5% (0.85 M) NaCl, 10 mM Na-phosphate, pH 8.0.

**Figure S10.** Weblogo representation of the alignment of internal sequences of *Hrr. lacusprofundi* FlaB1 and FlaB2 archaellins. In this representation, the overall height of a stack indicates the sequence conservation at that position, while the height of symbols within the stack indicates the relative frequency of each amino acid at that position (Crooks *et al.*, 2004). Left column: Amino acid residues 50-75; 53 *Halorubrum* FlaB1 sequences and 55 *Halorubrum* FlaB2 sequences were used. Right column: C-terminal sequences (∼ 25 amino acid residues from C-termini); 50 FlaB1 sequences and 53 FlaB2 sequences were used. The tables below show the corresponding characteristic signatures for FlaB1 and FlaB2. Conservative residues for both archaellins are colored red, conservative residues different for FlaB1 and FlaB2 are blue, and slightly conservative residues are black.

**Figure S11.** Schematic phylogenetic trees obtained using the Standard Protein BLAST https://blast.ncbi.nlm.nih.gov/Blast.cgi for selected archaellins (A), FlaG (B) and FlaH (C) of *Har. hispanica*, *Hbt.* Dl1, *Hbf. lacisalsi*, *Hfx. volcanii*, *Hpg. xanaduensis*, *Hrr. lacusprofundi*, *Hrr. saccharovorum*, *Nab. aegyptia* and *Nln. aegyptiacus*.

**Figure S12.** Mass spectrometry analysis of most prominent proteins in archaella isolated from wild type *Hrr. lacusprofundi* DL18 (top and middle) and ACAM 34 (bottom) strains. Protein coverage of archaellins B1 (WP_088901573.1) and B2 (WP_088901574.1/WP_015911241.1) are shown. In gray the full protein sequence is shown. In blue the unique peptides are depicted.

**Figure S13.** Mass spectrometry analysis of most prominent proteins in archaella isolated from *Hfx. volcanii* MT2 transformed with pAS5 (HL-B1B2-R). Protein coverage of archaellins B1 (WP_088901573.1) (top) and B2 (WP_088901574.1/WP_015911241.1) (bottom) are shown. In gray the full protein sequence is shown. In blue the unique peptides are depicted.

**Figure S14.** Mass spectrometry analysis of most prominent proteins in archaella isolated from *Hfx. volcanii* MT2 transformed with pAS6 (HL-B1-R) (top) and pAS7 (HL-B2-R) (bottom). Protein coverage of archaellins B1 (WP_088901573.1) and B2 (WP_088901574.1/WP_015911241.1) are shown. In gray the full protein sequence is shown. In blue the unique peptides are depicted. The red square corresponds to the possible formylation.

**Figure S15.** Mass spectrometry analysis of most prominent proteins in archaella isolated from wild type *Halorubrum saccharovorum* ATCC 29252. Protein coverage of archaellins B1 (WP_004047439.1) (top) and B2 (WP_004047440.1) (bottom) are shown. In gray the full protein sequence is shown. In blue the unique peptides are depicted.

**Figure S16.** Mass spectrometry analysis of most prominent proteins in archaella isolated from *Hfx. volcanii* MT2 transformed with pAS1 (HS-B1B2-R) (top and middle) and pAS2 (HS-B1-R) (bottom). Protein coverage of archaellins B1 (WP_004047439.1) and B2 (WP_004047440.1) are shown. In gray the full protein sequence is shown. In blue the unique peptides are depicted.

**Movie S1**. Swimming behavior of *Hfx. volcanii* MT2 transformed with pAS7 (HL-B2-R).

**Movie S2**. Swimming behavior of *Hfx. volcanii* MT2 transformed with pAS6 (HL-B1-R)

**Movie S3**. Swimming behavior of *Hfx. volcanii* MT2 transformed with pAS5 (HL-B1B2-R).

**Movie S4**. Swimming behavior of *Hfx. volcanii* MT2 transformed with with pMT21 (HV-A1A2),

