## Supplementary Figures 1-16 for "Interaction of two strongly divergent archaellins stabilizes the structure of the *Halorubrum* archaellum"

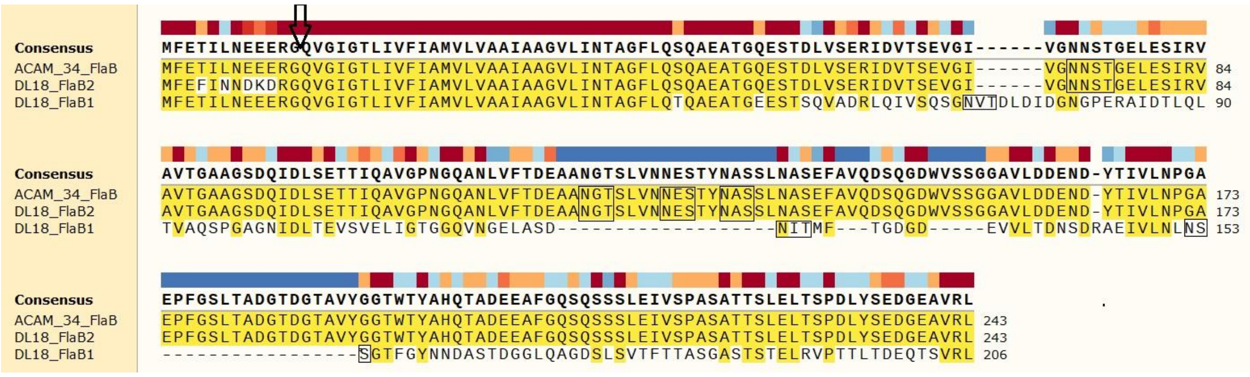
Consensus MFETILNEEERGQVGIGTLIVFIAMVLVAAIAAGVLINTAGFLQSQAEATGQESTDLVSERIDVTSEVGI------VGNNSTGELESIRV
ACAM_34_FlaB MFETILNEEERGQVGIGTLIVFIAMVLVAAIAAGVLINTAGFLQSQAEATGQESTDLVSERIDVTSEVGI------VGNNSTGELESIRV 84
DL18_FlaB2 MFEFINNDKDRGQVGIGTLIVFIAMVLVAAIAAGVLINTAGFLQSQAEATGQESTDLVSERIDVTSEVGI------VGNNSTGELESIRV 84
DL18_FlaB1 MFETILNEEERGQVGIGTLIVFIAMVLVAAIAAGVLINTAGFLQTQAEATGEESTSQVADRLQIVSQSGNVTDLDIDGNGPERAIDTLQL 90
Consensus AVTGAAGSDQIDLSETTIQAVGPNGQANLVFTDEAANGTSLVNNESTYNASSLNASEFAVQDSQGDWVSSGGAVLDDEND-YTIVLNPGA
ACAM_34_FlaB AVTGAAGSDQIDLSETTIQAVGPNGQANLVFTDEAANGTSLVNNESTYNASSLNASEFAVQDSQGDWVSSGGAVLDDEND-YTIVLNPGA 173
DL18_FlaB2 AVTGAAGSDQIDLSETTIQAVGPNGQANLVFTDEAANGTSLVNNESTYNASSLNASEFAVQDSQGDWVSSGGAVLDDEND-YTIVLNPGA 173
DL18_FlaB1 TVAQSPGAGNIDLTEVSVELIGTGGQVNGELASD-------------------NITMF---TGDGD-----EVVLTDNSDRAEIVLNLNS 153
Consensus EPFGSLTADGTDGTAVYGGTWTYAHQTADEEAFGQSQSSSLEIVSPASATTSLELTSPDLYSEDGEAVRL
ACAM_34_FlaB EPFGSLTADGTDGTAVYGGTWTYAHQTADEEAFGQSQSSSLEIVSPASATTSLELTSPDLYSEDGEAVRL 243
DL18_FlaB2 EPFGSLTADGTDGTAVYGGTWTYAHQTADEEAFGQSQSSSLEIVSPASATTSLELTSPDLYSEDGEAVRL 243
DL18_FlaB1 -----------------SGTFGYNNDASTDGGLQAGDSLSVTFTTASGASTSTELRVPTTLTDEQTSVRL 206

FS1

>NZ_MKFG01000003.1:83451-85227 Halorubrum lacusprofundi strain DL18 contig003, whole genome shotgun sequence

GGCGGGTCGGAACTCCGCGGAGCGGCCGAAGCGATCGGCGGCGGACCAGCGGATTCGGGAAGCGACTGTCCGACGATCGCACGCGAGAGCGTCGCGG
ATCCGCGTGCCGACCGCGCTCGATCCGACCGTCGATACGGAACTCTCACAGATCCAACGCCGATAATCCGATCACGACTTTGTTGATATATTCAAAA
ATCATTCGCTAGAAAATAATCCGATAATCGCAACACTGCAGCGTTTTATCAATTTGGATAATCTGCACGGAGTCTTTAATAGGGAAATTACACTTCG
TAGCGATAACGCCCGCCGGGCAGGCCGGGGGCCAACGCAACACACAATGTTCGAAACAATACTGAACGAGGAAGAGCGCGGTCAAGTGGGGATCGGG
ACCCTCATCGTGTTCATCGCGATGGTGCTGGTGGCGGCGATCGCCGCCGGCGTCCTGATCAACACGGCCGGCTTCCTTCAGACGCAGGCGGAAGCAA
CGGGCGAAGAGAGTACGAGTCAGGTGGCCGACCGGCTGCAGATCGTCAGCCAGTCGGGGAACGTGACTGATCTTGATATTGACGGAAACGGTCCCGA
ACGCGCGATCGATACCCTTCAGCTCACGGTCGCGCAGTCACCGGGTGCCGGAAACATCGATCTCACGGAAGTGAGCGTCGAACTGATCGGCACCGGT
GGGCAGGTCAACGGAGAACTGGCCAGCGATAACATCACGATGTTCACCGGAGACGGTGACGAAGTCGTTCTCACCGACAACAGCGACCGCGCTGAGA
TTGTGCTCAACCTCAATAGTAGCGGAACGTTCGGGTACAACAACGATGCAAGCACTGACGGCGGTCTACAGGCTGGCGACAGCCTCTCGGTGACGTT
CACGACCGCGTCCGGTGCATCGACGAGTACCGAACTGCGCGTCCCGACGACGCTGACTGACGAGCAAACCTCGGTGAGGCTGTAACAATGTTCGAGT
TTATCAACAACGACAAAGACCGCGGTCAGGTTGGCATCGGTACGCTCATCGTGTTCATCGCGATGGTACTGGTGGCGGCGATCGCCGCCGGTGTCCT
GATCAACACGGCCGGCTTCCTCCAGTCGCAGGCGGAAGCAACCGGACAGGAGAGTACGGATCTCGTCTCCGAACGGATCGACGTGACGAGCGAGGTC
GGTATCGTCGGGAACAACAGCACCGGCGAACTTGAGTCGATCCGCGTCGCCGTTACCGGTGCGGCCGGGTCCGATCAGATCGACTTATCAGAGACGA
CGATCCAAGCGGTCGGTCCGAACGGACAGGCGAACCTCGTGTTCACCGACGAGGCTGCAAATGGTACATCTCTAGTCAATAACGAGAGCACATACAA
TGCGAGCAGCCTCAATGCGAGTGAGTTCGCTGTGCAAGATTCCCAAGGCGATTGGGTCAGCAGTGGCGGTGCAGTGCTGGACGACGAGAACGATTAC
ACCATCGTCCTCAACCCTGGCGCAGAACCGTTCGGGAGCCTCACTGCGGACGGTACCGATGGCACAGCAGTCTACGGTGGAACGTGGACCTACGCTC
ACCAAACTGCAGACGAAGAGGCCTTCGGACAGAGCCAATCCTCGTCGCTCGAGATCGTCTCGCCCGCGTCGGCGACGACCTCACTCGAACTCACTTC
GCCCGACCTCTACAGCGAAGACGGCGAAGCGGTCCGGCTCTAACGAGCCTCGAATAGCCCCCTAACTACCTTCCCGGCGGATCCCTCCGCCCGGAAC
ACCTATTTTCTCGCACCACAACGGCTACGCC

>CP001365.1:2532506-2533659 Halorubrum lacusprofundi ATCC 49239 chromosome 1, complete sequence

GGCGGGTCGGAACTCCGCGGAGCGGCCGAAGCGATCGGCGGCGGACCAGCGGATTCGGGAAGCGACTGTCCGACGATCGCACGCGAGAGCGTCGCGG
ATCCGCGTGCCGACCGCGCTCGATCCGACCGTCGATACGGAACTCTCACAGATCCAACGCCGATAATCCGATCACGACTTTGTTGATATATTCAAAA
ATCATTCGCTAGAAAATAATCCGATAATCGCAACACTGCAGCGTTTTATCAATTTGGATAATCTGCACGGAGTCTTTAATAGGGAAATTACACTTCG
TAGCGATAACGCCCGCCGGGCAGGCCGGGGGCCAACGCAACACACAATGTTCGAAACAATACTGAACGAGGAAGAGCGCGGTCAGGTTGGCATCGGT
ACGCTCATCGTGTTCATCGCGATGGTACTGGTGGCGGCGATCGCCGCCGGTGTCCTGATCAACACGGCCGGCTTCCTCCAGTCGCAGGCGGAAGCAA
CCGGACAGGAGAGTACGGATCTCGTCTCCGAACGGATCGACGTGACGAGCGAGGTCGGTATCGTCGGGAACAACAGCACCGGCGAACTTGAGTCGAT
CCGCGTCGCCGTTACCGGTGCGGCCGGGTCCGATCAGATCGACTTATCAGAGACGACGATCCAAGCGGTCGGTCCGAACGGACAGGCGAACCTCGTG
TTCACCGACGAGGCTGCAAATGGTACATCTCTAGTCAATAACGAGAGCACATACAATGCGAGCAGCCTCAATGCGAGTGAGTTCGCTGTGCAAGATT
CCCAAGGCGATTGGGTCAGCAGTGGCGGTGCAGTGCTGGACGACGAGAACGATTACACCATCGTCCTCAACCCTGGCGCAGAACCGTTCGGGAGCCT
CACTGCGGACGGTACCGATGGCACAGCAGTCTACGGTGGAACGTGGACCTACGCTCACCAAACTGCAGACGAAGAGGCCTTCGGACAGAGCCAATCC
TCGTCGCTCGAGATCGTCTCGCCCGCGTCGGCGACGACCTCACTCGAACTCACTTCGCCCGACCTCTACAGCGAAGACGGCGAAGCGGTCCGGCTCT
AACGAGCCTCGAATAGCCCCCTAACTACCTTCCCGGCGGATCCCTCCGCCCGGAACACCTATTTTCTCGCACCACAACGGCTACGCC

FS2

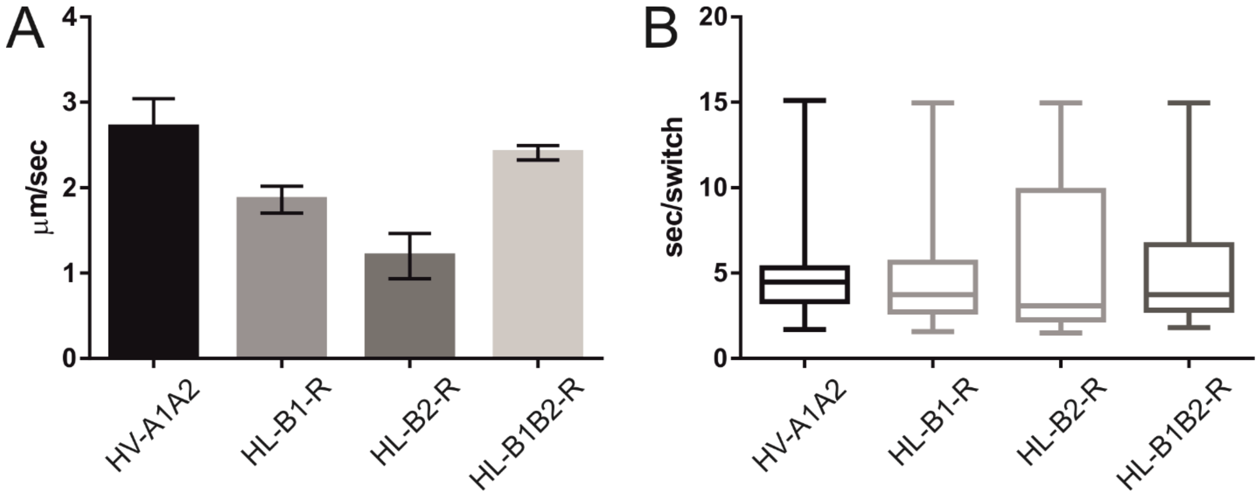
A
4
3
2
1
0
μm/sec
HV-A1A2
HL-B1-R
HL-B2-R
HL-B1B2-R
B
20
15
10
5
0
sec/switch
HV-A1A2
HL-B1-R
HL-B2-R
HL-B1B2-R

FS3

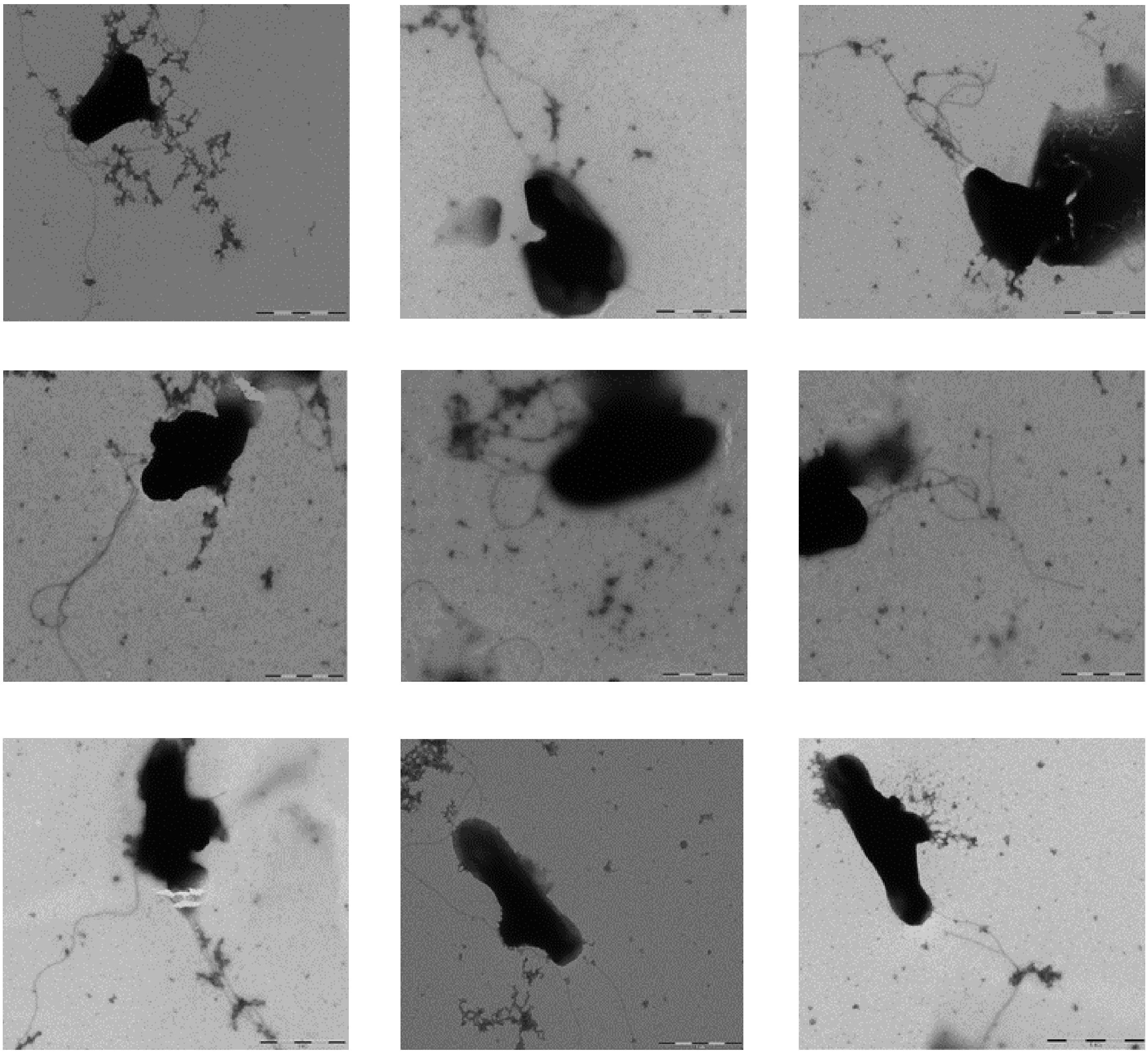

FS4

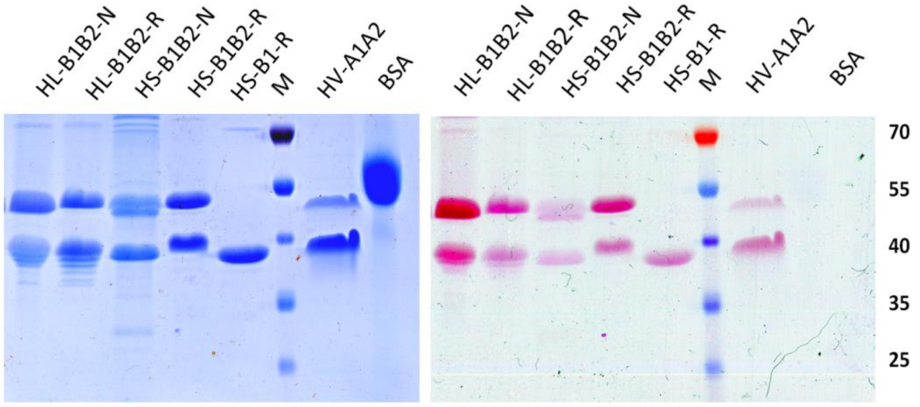
HL-B1B2-N
HL-B1B2-R
HS-B1B2-N
HS-B1B2-R
HS-B1-R
M
HV-A1A2
BSA
HL-B1B2-N
HL-B1B2-R
HS-B1B2-N
HS-B1B2-R
HS-B1-R
M
HV-A1A2
BSA
70
55
40
35
25

FS5

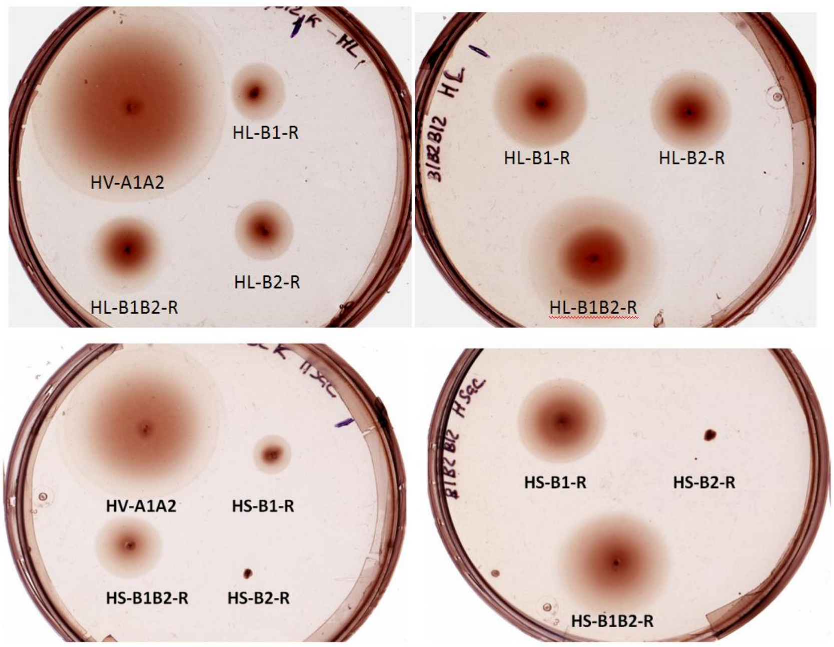
HV-A1A2
HL-B1-R
HL-B1B2-R
HL-B2-R
HL-B1-R
HL-B2-R
HL-B1B2-R
HV-A1A2
HS-B1-R
HS-B1B2-R
HS-B2-R
HS-B1-R
HS-B2-R
HS-B1B2-R

FS6

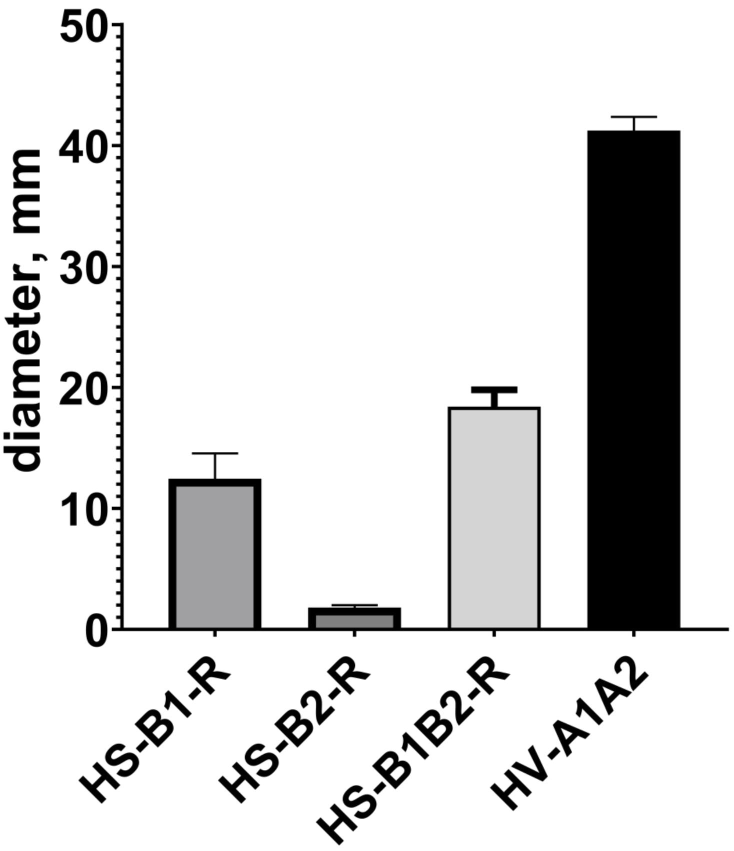


FS7

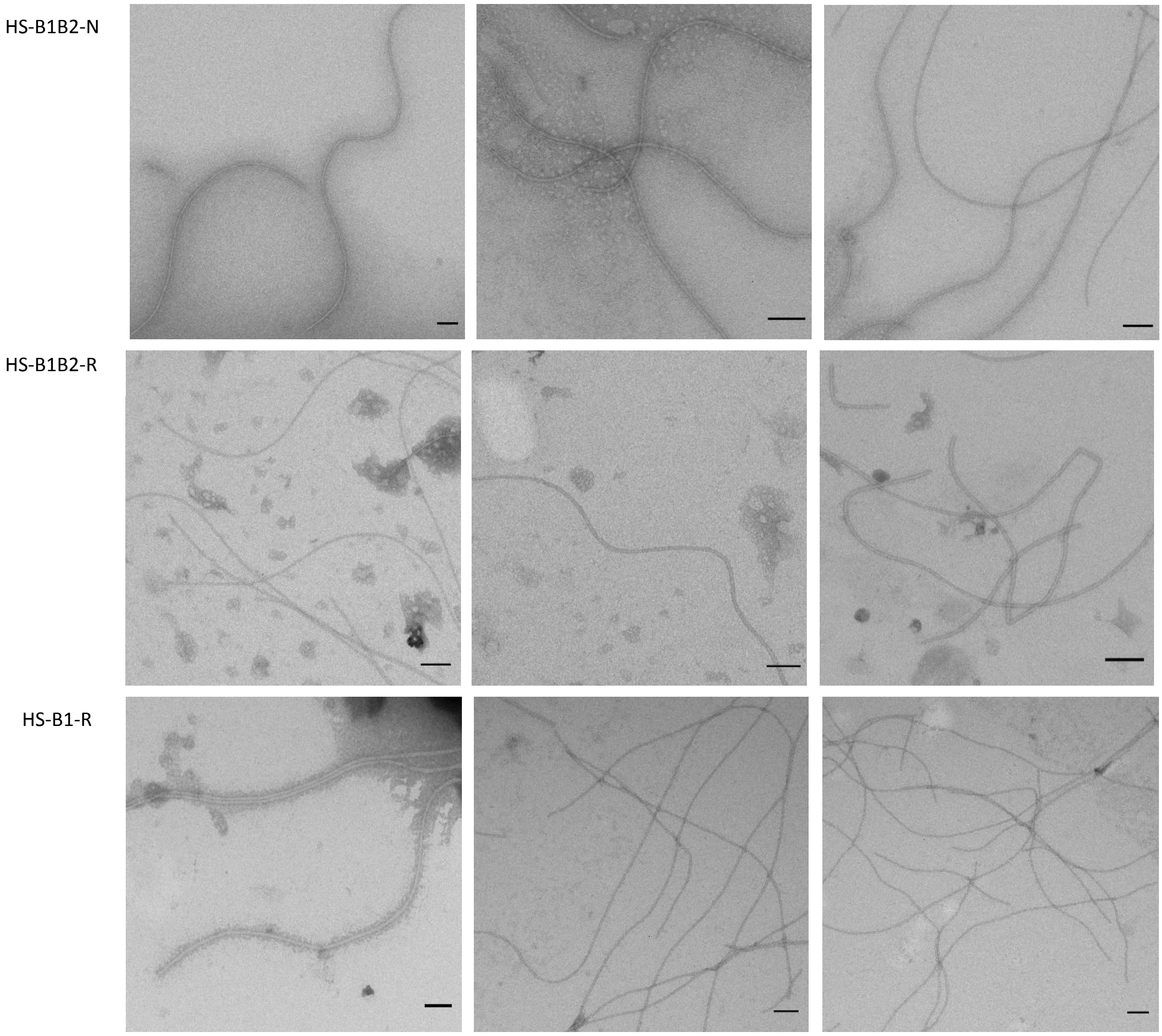


FS8

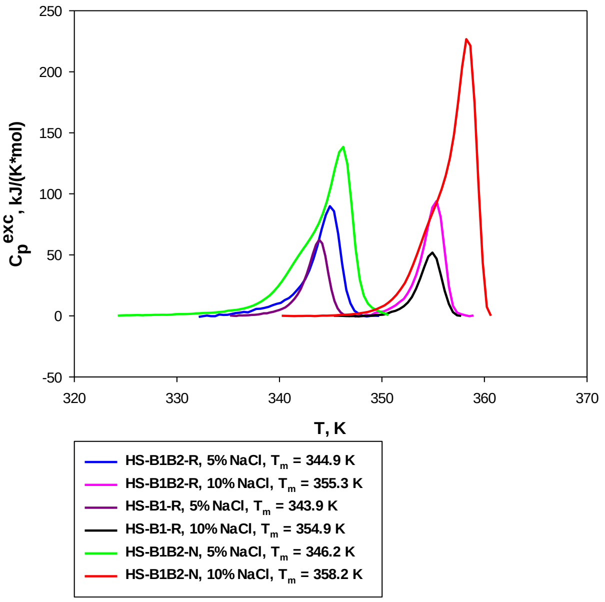

250
200
150
100
50
0
-50
320
330
340
350
360
370
Cp exc, kJ/(K*mol)
T, K
HS-B1B2-R, 5% NaCl, Tm = 344.9 K
HS-B1B2-R, 10% NaCl, Tm = 355.3 K
HS-B1-R, 5% NaCl, Tm = 343.9 K
HS-B1-R, 10% NaCl, Tm = 354.9 K
HS-B1B2-N, 5% NaCl, Tm = 346.2 K
HS-B1B2-N, 10% NaCl, Tm = 358.2 K


FS9

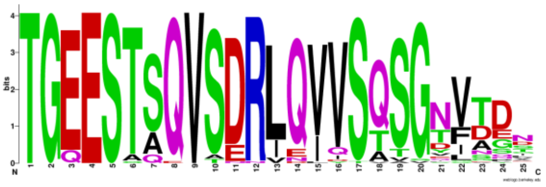

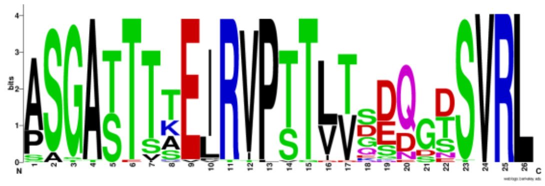

FlaB1

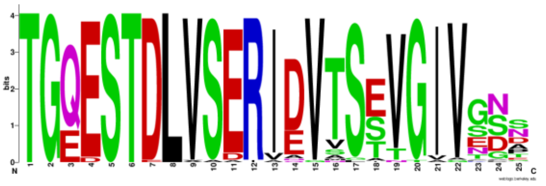

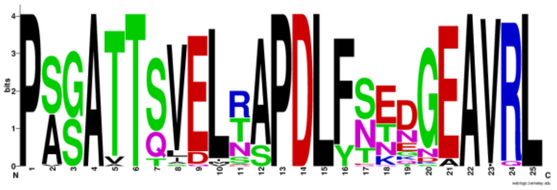

FlaB2

| | 54 | 55 | 56 | 57 | 58 | 59 | 60 | 61 | 62 | 63 | 64 | 65 | 66 | 66 | 67 | 68 |
|---|---|---|---|---|---|---|---|---|---|---|---|---|---|---|---|---|
| FlaB1 | S | T | S[A] | Q | V | S | D[E] | R | L[I] | Q | V[I] | V | S | Q[TA] | S | G |
| FlaB2 | S | T | D | L | V | S | E[D] | R | I[V] | D[E] | V | T | S | E[ST] | V | G |

| | 17 | 16 | 15 | 14 | 13 | 12 | 11 | 10 | 9 | 8 | 7 | 6 | 5 | 4 | 3 | 2 | 1 |
|---|---|---|---|---|---|---|---|---|---|---|---|---|---|---|---|---|---|
| FlaB1 | E | I[L] | R | V | P | T[S] | T | L[V] | T[V] | D[E] | Q[D] | G[D] | D[TS] | S | V | R | L |
| FlaB2 | E | L | R[TN] | A[S] | P | D | L | F[Y] | S[N] | E[TN] | D[NE] | G | E | A | V | R | L |

FS10

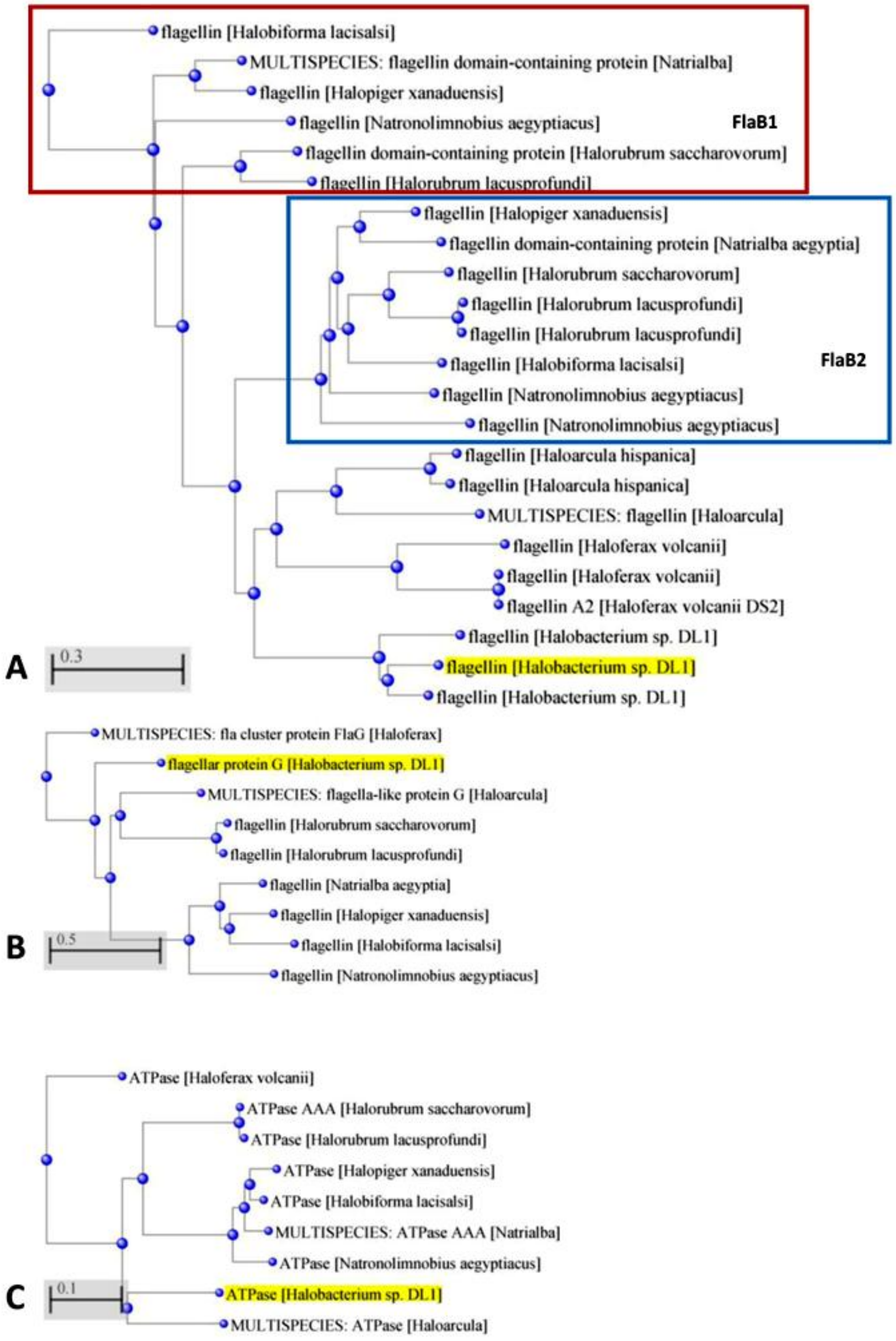
flagellin [Halobiforma lacisalsi]
MULTISPECIES: flagellin domain-containing protein [Natrialba]
flagellin [Halopiger xanaduensis]
flagellin [Natronolimnobius aegyptiacus]
FlaB1
flagellin domain-containing protein [Halorubrum saccharovorum]
flagellin [Halorubrum lacusprofundi]
flagellin [Halopiger xanaduensis]
flagellin domain-containing protein [Natrialba aegyptia]
flagellin [Halorubrum saccharovorum]
flagellin [Halorubrum lacusprofundi]
flagellin [Halorubrum lacusprofundi]
flagellin [Halobiforma lacisalsi]
FlaB2
flagellin [Natronolimnobius aegyptiacus]
flagellin [Natronolimnobius aegyptiacus]
flagellin [Haloarcula hispanica]
flagellin [Haloarcula hispanica]
MULTISPECIES: flagellin [Haloarcula]
flagellin [Haloferax volcanii]
flagellin [Haloferax volcanii]
flagellin A2 [Haloferax volcanii DS2]
flagellin [Halobacterium sp. DL1]
A
0.3
flagellin [Halobacterium sp. DL1]
flagellin [Halobacterium sp. DL1]
MULTISPECIES: fla cluster protein FlaG [Haloferax]
flagellar protein G [Halobacterium sp. DL1]
MULTISPECIES: flagella-like protein G [Haloarcula]
flagellin [Halorubrum saccharovorum]
flagellin [Halorubrum lacusprofundi]
flagellin [Natrialba aegyptia]
flagellin [Halopiger xanaduensis]
B
0.5
flagellin [Halobiforma lacisalsi]
flagellin [Natronolimnobius aegyptiacus]
ATPase [Haloferax volcanii]
ATPase AAA [Halorubrum saccharovorum]
ATPase [Halorubrum lacusprofundi]
ATPase [Halopiger xanaduensis]
ATPase [Halobiforma lacisalsi]
MULTISPECIES: ATPase AAA [Natrialba]
ATPase [Natronolimnobius aegyptiacus]
C
0.1
ATPase [Halobacterium sp. DL1]
MULTISPECIES: ATPase [Haloarcula]

FS11

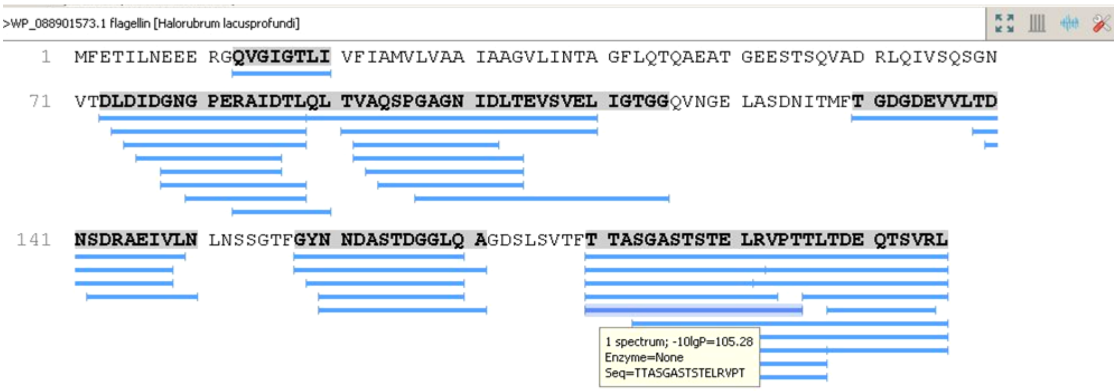
>WP_088901573.1 flagellin [Halorubrum lacusprofundi]
1 MFETILNEEE RGQVGIGTLI VFIAMVLVAA IAAGVLINTA GFLQTQAEAT GEESTSQVAD RLQIVSQSGN
71 VTDLDIDGNG PERAIDTLQL TVAQSPGAGN IDLTEVSVEL IGTGGQVNGE LASDNITMFT GDGDEVVLTD
141 NSDRAEIVLN LNSSGTFGYN NDASTDGGLQ AGDSLSVTFT TASGASTSTE LRVPTTLTDE QTSVRL
1 spectrum; -10lgP=105.28
Enzyme=None
Seq=TTASGASTSTELRVPT

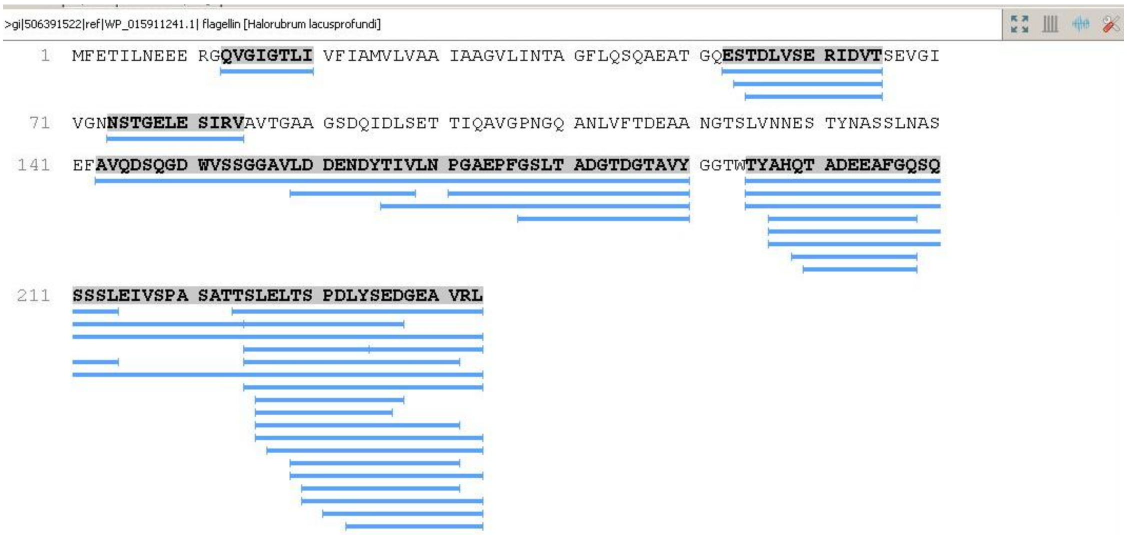
>gi|506391522|ref|WP_015911241.1| flagellin [Halorubrum lacusprofundi]
1 MFETILNEEE RGQVGIGTLI VFIAMVLVAA IAAGVLINTA GFLQSQAEAT GQESTDLVSE RIDVTSEVGI
71 VGNNSTGELE SIRVAVTGAA GSDQIDLSET TIQAVGPNGQ ANLVFTDEAA NGTSLVNNES TYNASSLNAS
141 EFAVQDSQGD WVSSGGAVLD DENDYTIVLN PGAEPFGSLT ADGTDGTAVY GGTWTYAHQT ADEEAFGQSQ
211 SSSLEIVSPA SATTSLELTS PDLYSEDGEA VRL

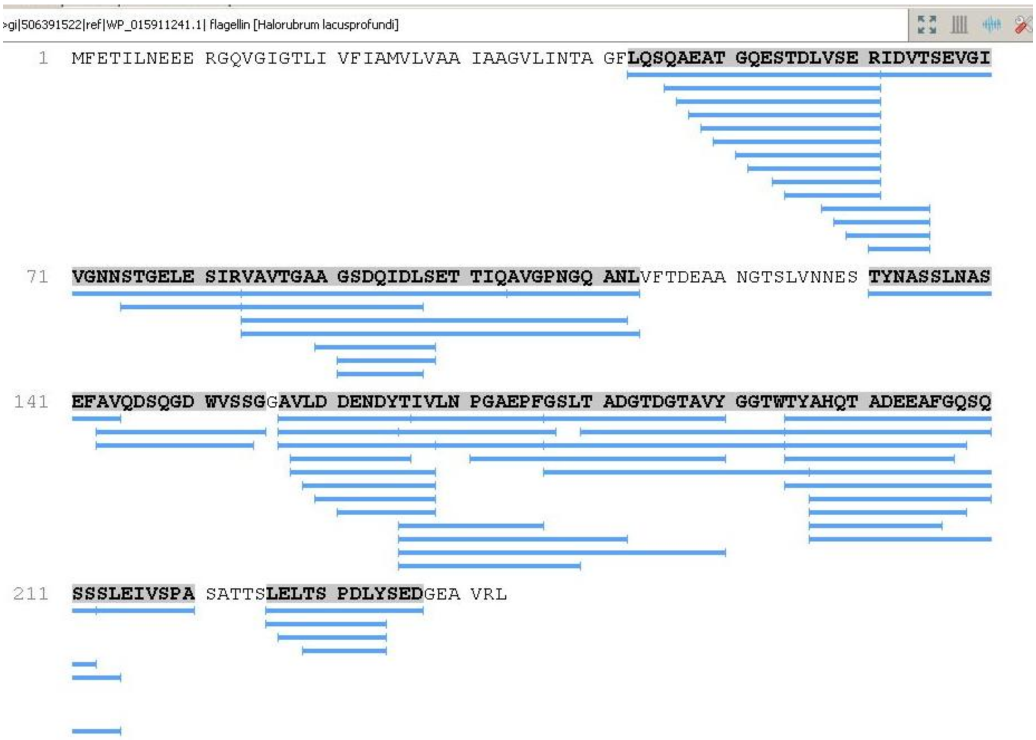
>gi|506391522|ref|WP_015911241.1| flagellin [Halorubrum lacusprofundi]
1 MFETILNEEE RGQVGIGTLI VFIAMVLVAA IAAGVLINTA GFLQSQAEAT GQESTDLVSE RIDVTSEVGI
71 VGNNSTGELE SIRVAVTGAA GSDQIDLSET TIQAVGPNGQ ANLVFTDEAA NGTSLVNNES TYNASSLNAS
141 EFAVQDSQGD WVSSGGAVLD DENDYTIVLN PGAEPFGSLT ADGTDGTAVY GGTWTYAHQT ADEEAFGQSQ
211 SSSLEIVSPA SATTSLELTS PDLYSEDGEA VRL

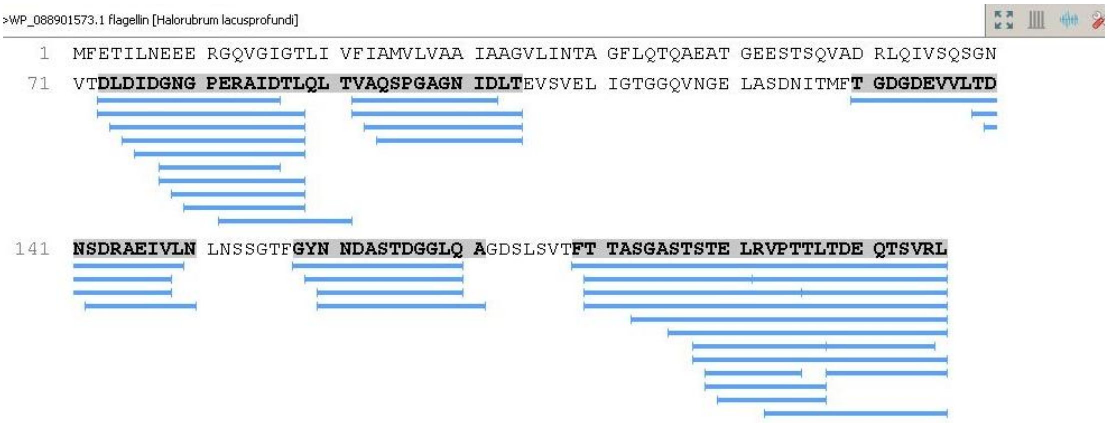
>WP_088901573.1 flagellin [Halorubrum lacusprofundi]
1 MFETILNEEE RGQVGIGTLI VFIAMVLVAA IAAGVLINTA GFLQTQAEAT GEESTSQVAD RLQIVSQSGN
71 VTDLDIDGNG PERAIDTLQL TVAQSPGAGN IDLTEVSVEL IGTGGQVNGE LASDNITMFT GDGDEVVLTD
141 NSDRAEIVLN LNSSGTFGYN NDASTDGGLQ AGDSLSVTFT TASGASTSTE LRVPTTLTDE QTSVRL

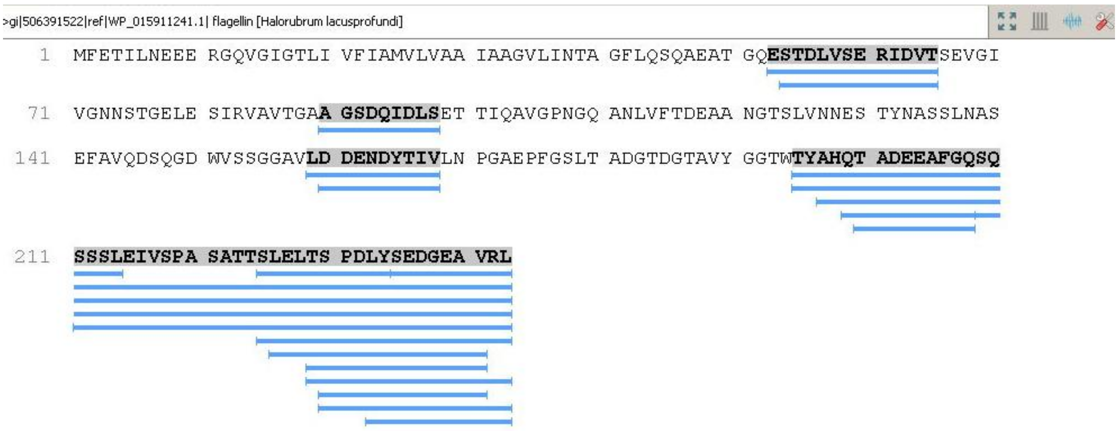
>gi|506391522|ref|WP_015911241.1| flagellin [Halorubrum lacusprofundi]
1 MFETILNEEE RGQVGIGTLI VFIAMVLVAA IAAGVLINTA GFLQSQAEAT GQESTDLVSE RIDVTSEVGI
71 VGNNSTGELE SIRVAVTGAA GSDQIDLSET TIQAVGPNGQ ANLVFTDEAA NGTSLVNNES TYNASSLNAS
141 EFAVQDSQGD WVSSGGAVLD DENDYTIVLN PGAEPFGSLT ADGTDGTAVY GGTWTYAHQT ADEEAFGQSQ
211 SSSLEIVSPA SATTSLELTS PDLYSEDGEA VRL

FS13

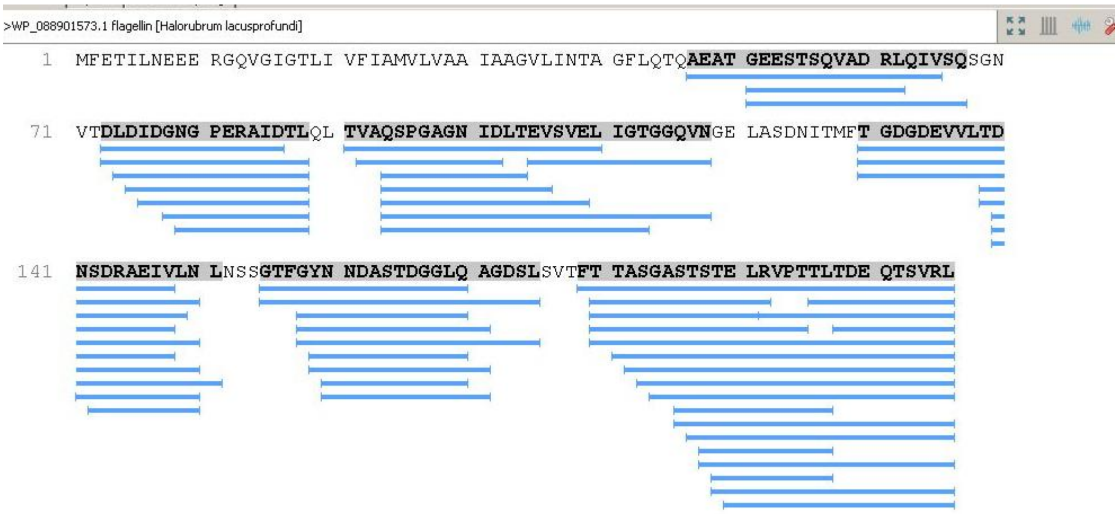

>WP_088901573.1 flagellin [Halorubrum lacusprofundi]
1 MFETILNEEE RGQVGIGTLI VFIAMVLVAA IAAGVLINTA GFLQTQAEAT GEESTSQVAD RLQIVSQSGN
71 VTDLDIDGNG PERAIDTLQL TVAQSPGAGN IDLTEVSVEL IGTGGQVNGE LASDNITMFT GDGDEVVLTD
141 NSDRAEIVLN LNSSGTFGYN NDASTDGGLQ AGDSLSVTFT TASGASTSTE LRVPTTLTDE QTSVRL


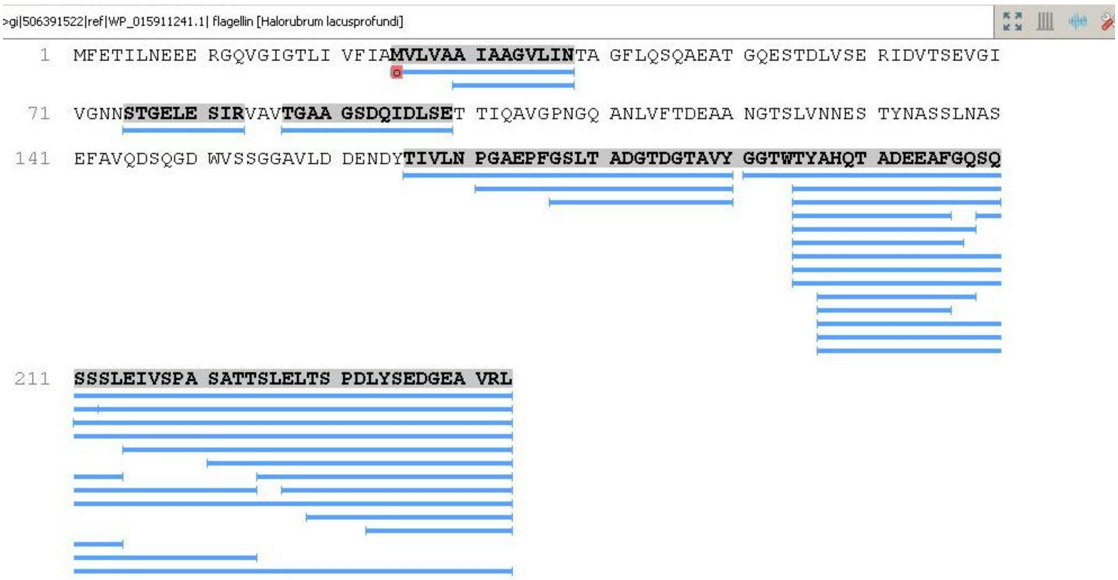

>gi|506391522|ref|WP_015911241.1| flagellin [Halorubrum lacusprofundi]
1 MFETILNEEE RGQVGIGTLI VFIAMVLVAA IAAGVLINTA GFLQSQAEAT GQESTDLVSE RIDVTSEVGI
71 VGNNSTGELE SIRVAVTGAA GSDQIDLSET TIQAVGPNGQ ANLVFTDEAA NGTSLVNNES TYNASSLNAS
141 EFAVQDSQGD WVSSGGAVLD DENDYTIVLN PGAEPFGSLT ADGTDGTAVY GGTWTYAHQT ADEEAFGQSQ
211 SSSLEIVSPA SATTSLELTS PDLYSEDGEA VRL


FS14

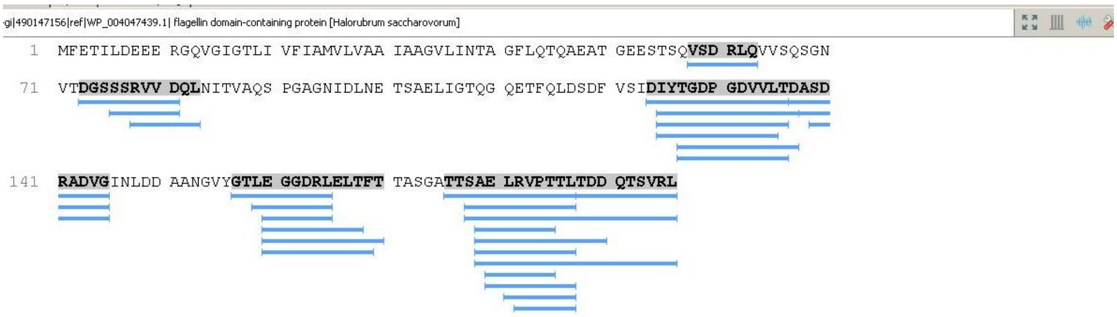
gi|490147156|ref|WP_004047439.1| flagellin domain-containing protein [Halorubrum saccharovorum]
1 MFETILDEEE RGQVGIGTLI VFIAMVLVAA IAAGVLINTA GFLQTQAEAT GEESTSQVSD RLQVVSQSGN
71 VTDGSSSRVV DQLNITVAQS PGAGNIDLNE TSAELIGTQG QETFQLDSDF VSIDIYTGDP GDVVLTDASD
141 RADVGINLDD AANGVYGTLE GGDRLELTFT TASGATTSAE LRVPTTLTDD QTSVRL

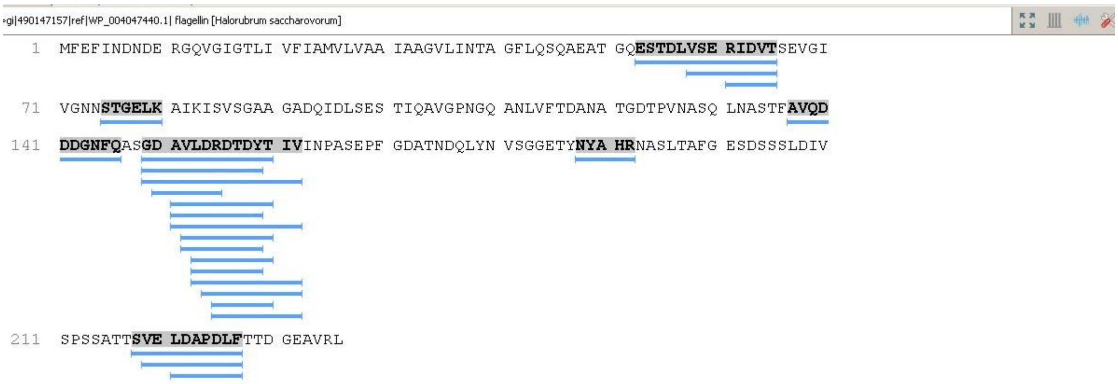
gi|490147157|ref|WP_004047440.1| flagellin [Halorubrum saccharovorum]
1 MFEFINDNDE RGQVGIGTLI VFIAMVLVAA IAAGVLINTA GFLQSQAEAT GQESTDLVSE RIDVTSEVGI
71 VGNNSTGELK AIKISVSGAA GADQIDLSES TIQAVGPNGQ ANLVFTDANA TGDTPVNASQ LNASTFAVQD
141 DDGNFQASGD AVLDRDTDYT IVINPASEPF GDATNDQLYN VSGGETYNYA HRNASLTAFG ESDSSSLDIV
211 SPSSATTSVE LDAPDLFTTD GEAVRL

FS15

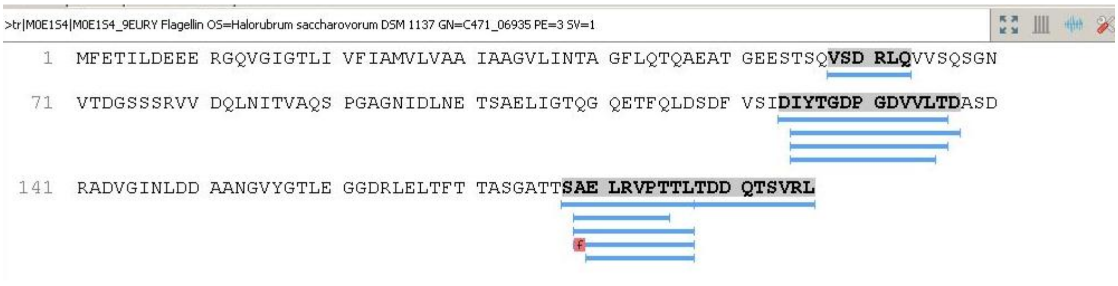
>tr|M0E1S4|M0E1S4_9EURY Flagellin OS=Halorubrum saccharovorum DSM 1137 GN=C471_06935 PE=3 SV=1
1 MFETILDEEE RGQVGIGTLI VFIAMVLVAA IAAGVLINTA GFLQTQAEAT GEESTSQVSD RLQVVSQSGN
71 VTDGSSSRVV DQLNITVAQS PGAGNIDLNE TSAELIGTQG QETFQLDSDF VSIDIYTGDP GDVVLTDASD
141 RADVGINLDD AANGVYGTLE GGDRLELTFT TASGATTSAE LRVPTTLTDD QTSVRL

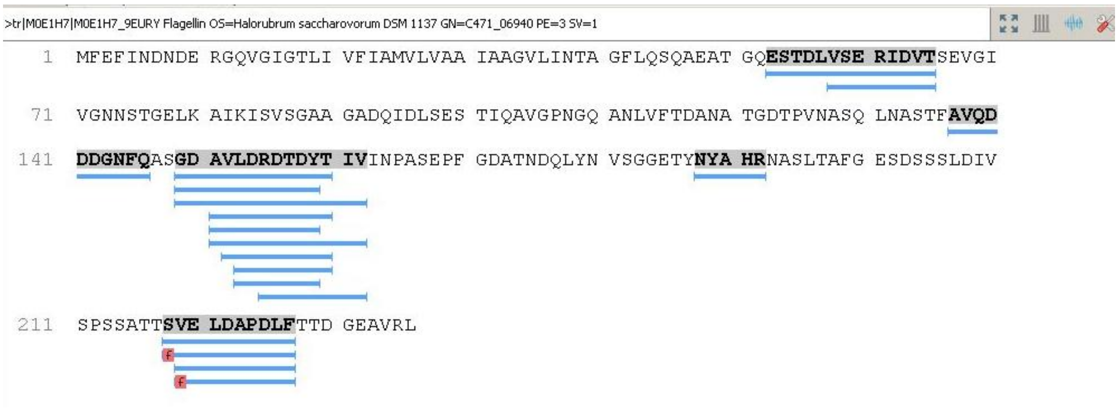
>tr|M0E1H7|M0E1H7_9EURY Flagellin OS=Halorubrum saccharovorum DSM 1137 GN=C471_06940 PE=3 SV=1
1 MFEFINDNDE RGQVGIGTLI VFIAMVLVAA IAAGVLINTA GFLQSQAEAT GQESTDLVSE RIDVTSEVGI
71 VGNNSTGELK AIKISVSGAA GADQIDLSES TIQAVGPNGQ ANLVFTDANA TGDTPVNASQ LNASTFAVQD
141 DDGNFQASGD AVLDRDTDYT IVINPASEPF GDATNDQLYN VSGGETYNYA HRNASLTAFG ESDSSSLDIV
211 SPSSATTSVE LDAPDLFTTD GEAVRL

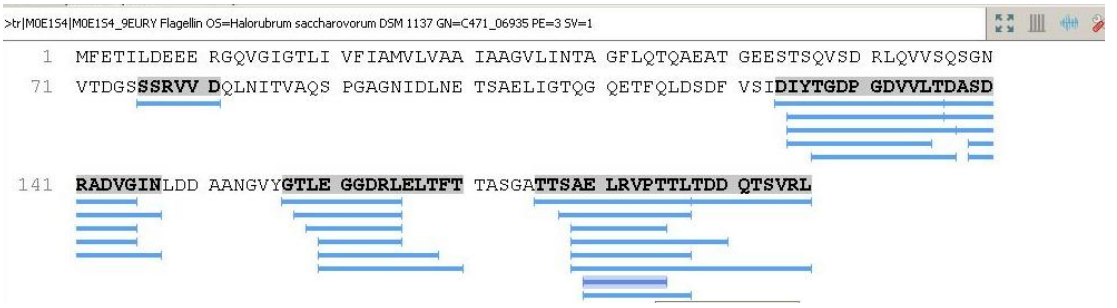
>tr|M0E1S4|M0E1S4_9EURY Flagellin OS=Halorubrum saccharovorum DSM 1137 GN=C471_06935 PE=3 SV=1
1 MFETILDEEE RGQVGIGTLI VFIAMVLVAA IAAGVLINTA GFLQTQAEAT GEESTSQVSD RLQVVSQSGN
71 VTDGSSSRVV DQLNITVAQS PGAGNIDLNE TSAELIGTQG QETFQLDSDF VSIDIYTGDP GDVVLTDASD
141 RADVGINLDD AANGVYGTLE GGDRLELTFT TASGATTSAE LRVPTTLTDD QTSVRL

FS16
